# Powering methanogenesis from fatty acids by a twin-heme-mediated reverse redox-loop

**DOI:** 10.64898/2026.08.18.745454

**Authors:** Dennis Kosian, Lin Zhang, Lena Appel, Lorenz Heidinger, Friedel Drepper, Pitter Huesgen, Oliver Einsle, Matthias Boll

## Abstract

The conversion of organic matter into methane is central to the global carbon cycle and engineered biogas production. In this process, syntrophic bacteria oxidize fatty acid fermentation products to acetate, coupled to the generation of H_2_ or formate, which are subsequently utilized by methanogenic archaea. During fatty acid β-oxidation, a membrane-bound electron-transferring flavoprotein (ETF):methylmenaquinone (MMK) oxidoreductase complex (EMO) has been proposed to drive endergonic electron transfer from reduced ETF to CO_2_ through a reverse redox loop, with its mechanistic basis remaining unresolved. Here we report cryo-electron microscopy structures of EMO and the EMO-ETF complex from *Syntrophus aciditrophicus* at 2.0 and 3.0 Å resolution, respectively. Complex formation induces substantial conformational rearrangements in ETF, positioning its flavin for efficient electron transfer to non-cubane [4Fe:4S] and [4Fe:5S] clusters. The membrane-integral domain harbors three heme *b* cofactors, including a specialized twin-heme unit that mediates proton-motive-force-driven MMK reduction. The structural and functional similarity of EMOs to heterodisulfide reductases, together with their broad distribution across bacteria and archaea, suggests an evolutionary link between methanogenesis and fatty acid β-oxidation, illustrating how ancient redox systems were repurposed for new metabolic functions.

## Introduction

The microbial conversion of biomass into methane is a central process in the global carbon cycle accounting for nearly half of the methane produced worldwide^1,2^. It also underpins sustainable biogas generation from waste or wastewater in engineered systems such as sewage treatment and biogas plants^3,4^. Methanogenic biomass degradation proceeds through three main stages: hydrolysis and primary fermentation of biomass to short-chain alcohols and fatty acids; oxidation of these intermediates to acetate coupled to H^+^ or CO_2_ reduction; and methanogenesis, in which archaea convert CO_2_, formate, acetate or methylated compounds into methane^5^. The latter stages rely on interspecies electron transfer between fermenters and methanogens, mediated either by diffusible carriers such as H_2_ or formate^6,7^, or by direct electron transfer through conductive pili^8^. More recently, formate disproportionation to CO_2_ and methanol has been reported, providing evidence for syntrophic methylotrophic methanogenesis^9^.

The oxidation of short-chain fatty acids to acetate coupled to H^+^ or CO_2_ reduction is endergonic under standard conditions (e.g., butyrate → 2 acetate + 2 H_2_; Δ*G*^0^’ = +48 kJ*·*mol^−1^) but becomes favorable when H_2_ or formate are kept below ~10 Pa or 10 µM, yielding *E*’ ≈ –290 mV^5,10^. In syntrophic deltaproteobacteria such as *Syntrophus aciditrophicus*, butyrate oxidation occurs in two steps with distinct redox potentials, catalysed by electron transferring flavoprotein (ETF)-dependent butyryl-CoA dehydrogenase (DH, *E*^0^’ ≈ –10 mV) and NAD^+^-dependent 3-hydroxybutyryl-CoA DH (*E*^0^’ ≈ –250 mV)^11,12^. Whereas cytoplasmic formate dehydrogenase (FDH) can regenerate NAD^+^ by reduction of CO_2_ under low formate concentrations, reduced ETF is insufficient to drive CO_2_ reduction (Δ*G*^0^’ > 50 kJ*·*mol^−1^) even under syntrophic conditions^13^.

This long-standing energetic challenge of syntrophic methane formation was recently elucidated by the identification of methylmenaquinone (MMK), a membrane-bound electron carrier (*E*^0^’ = –156 mV) that serves as substrate for membrane-bound ETF:MMK oxidoreductase in *S. aciditrophicus* (*Sa*EMO)^14^. The unusual juxtaposition of *Sa*EMO toward the cytoplasm and membrane-bound FDH (mFDH) toward the periplasm suggested that the endergonic reduction of CO_2_ by ETF_red_ may be driven by a reverse redox loop powered by the proton-motive force (pmf) (Fig. 1). ETF-derived electrons typically reduce quinones on the cytoplasmic side of the membrane, coupled to proton uptake, whereas quinol oxidation on the periplasmic side releases protons to generate the transmembrane proton gradient. Reversing this process in *Sa*EMO implies that periplasmic proton uptake is exergonic and covers the energy differential between the electrons delivered by ETF and the acceptor MMK. Exploiting the proton-motive force to lower the midpoint potential of electrons is an unusual but not unprecedented mechanism that was also reported for the RNF complex of *Azotobacter vinelandii*^15^. Pmf is generated by proton-pumping ATPase fuelled by ATP-forming acetyl-CoA synthetase^16^. *Sa*EMO’s cytoplasmic domain is suggested to bind [4Fe:4S] clusters and resembles subunits of membrane-bound (HdrD) or soluble (HdrBC) heterodisulfide reductases (HDRs) from methanogens^17^. Its transmembrane (TM) domain resembles that of HdrE in membrane-bound HDRs and is proposed to bind two heme *b* with markedly differing redox potentials (*E*^0^’ = –80 mV and *E*^0^’ = –220 mV). The resulting 140 mV redox barrier is thought to be overcome by the periplasmic consumption and cytoplasmic release of protons during the reduction of CO_2_ by ETF_red_ and MMK_ox/red_ (Fig. 1)^14^. This reverse redox loop can also operate in the forward direction during growth on crotonate without a methanogenic partner. In the process of enoyl-CoA CoA respiration, the formate generated from CO_2_ during crotonyl-CoA oxidation is used to reduce a second crotonyl-CoA to butyryl-CoA via EMO and ETF^18^.

**Figure 1.**
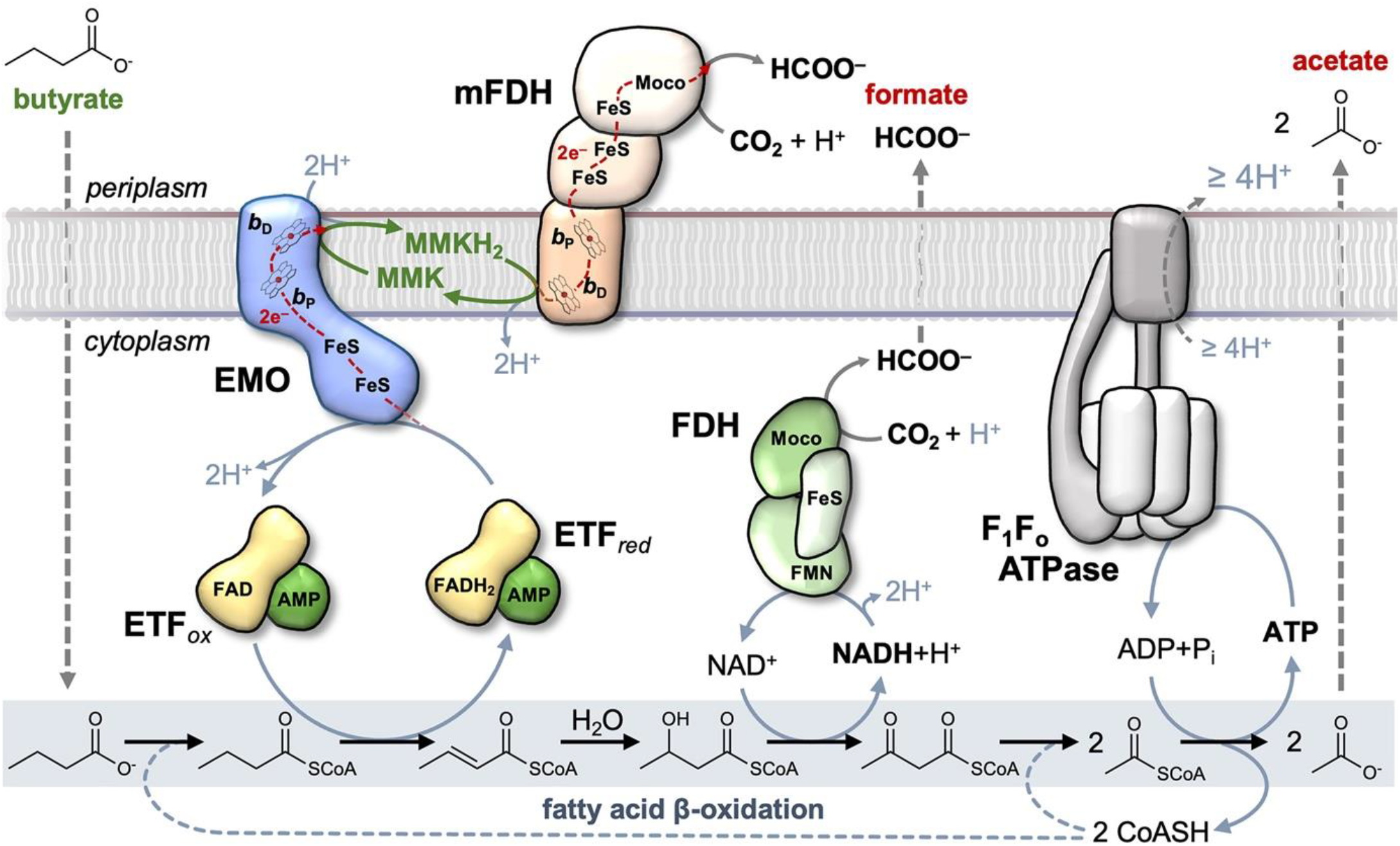
The pathway of syntrophic butyrate oxidation coupled to CO_2_ reduction. Electron transfer from NADH to CO_2_ is feasible under syntrophic conditions, whereas transfer from reduced ETF_red_ to CO_2_ must be driven by a MMK-dependent reverse redox loop formed by EMO and mFDH. The proton motive force powering this process is generated by the F_o_F_1_ ATP synthase running in reverse to hydrolyse ATP produced via substrate-level phosphorylation from acetyl-CoA. H^+^ transport associated with substrate import and product export is not shown.

Phylogenetic analyses revealed that EMO occurs in all micro-organisms containing MMK or menaquinone (MK) that can use fatty acids as growth substrates^14^. A single copy of a putative *emo* gene was also identified in *Mycobacterium tuberculosis*, whose persistence depends on the β-oxidation of host lipids^19^. Deleting *emo* in *M. tuberculosis* (also referred to as *etfD*) abolished growth on fatty acids and cholesterol, while growth on glycerol remained unaffected^20^. A recent 3.2 Å resolution cryoelectron microscopy (cryo-EM) structure of EMO (EtfD) from *M. smegmatis* (*Ms*EMO) revealed four FeS clusters in the cytoplasmic domains, including two canonical [4Fe:4S] clusters, a non-cubane [4Fe:4S] and a linear [3Fe:4S] cluster^21^. Non-cubane [4Fe:4S] clusters were previously identified in the HdrB subunit of soluble HDRs and proposed for homologous membrane-bound HdrD subunits in methanogens^22,23^, where they form catalytic sites for heterodisulfide reduction. In EMOs, however, they appear to function in electron transfer. Unlike *Sa*EMO, *Ms*EMO contains only a single heme *b*^21^, consistent with electron transfer to MK without energetic coupling.

To resolve the structural and mechanistic basis of the key step of methanogenesis from fatty acids, we determined cryo-EM structures of *Sa*EMO and the *Sa*EMO-*Sa*EtfAB complex at resolutions of 2.0 Å and 3.0 Å, respectively complemented by biochemical and spectroscopic analyses. Our results uncover unprecedented non-cubane [4Fe:5S] and twin-heme cofactors that enable a pmf-driven reverse redox loop to support the otherwise highly endergonic CO_2_ reduction by ETF_red_. The striking architectural and mechanistic parallels between EMOs and HDRs suggest an evolutionary link between methanogenesis and fatty acid β-oxidation.

## Results and Discussion

### A three-domain architecture with non-cubane FeS clusters and stacked twin-hemes *b*

To elucidate the structural basis of the *Sa*EMO-mediated electron transfer, both the EtfAB and *Sa*EMO components were enriched from *S. aciditrophicus* wild-type cells as described previously^14^. Purified *Sa*EMO migrated as a single band at approximately 80 kDa in SDS-PAGE (Supplementary Fig. 1a) and exhibited the characteristic UV/vis spectra of the oxidized (as isolated) and reduced states, which were dominated by the absorption of the heme *b* cofactors and, to a lesser extent, by the FeS clusters (Supplementary Fig 1b). The functional integrity of *Sa*EMO was verified by monitoring the time-dependent reduction of its heme *b* cofactors in the presence of EtfAB using cyclohexanoyl-CoA along with cyclohexanoyl-CoA dehydrogenase^24^ as an electron donor system (Supplementary Fig. 1c)^14^.

Single-particle cryo-EM data for *Sa*EMO was used to generate a 3D reconstruction from 267,857 individual particles to an overall resolution of 2.0 Å (Fig. 2a, Supplementary Table 1 and Supplementary Fig. 2). An atomic model built into this map comprised residues V2 to P695, lacking only the N-terminal methionine and nine residues at the C-terminus of the 706 amino acid protein. *Sa*EMO is an integral membrane protein comprising a membrane domain with five transmembrane (TM) helices that binds the heme *b* cofactors, and two iron-sulfur cluster binding domains in the cytoplasm (Fig. 2b). While the C-terminus is in the cytoplasm, the periplasmic N-terminus contains an unprocessed, inverted signal sequence that forms the first TM helix of the membrane-spanning region, which extends to residue 260. The TM domain of *Sa*EMO is homologous in sequence and structure to proteins of the NarI family, such as the NarI subunit of nitrate reductase A (NarGHI, PDB 1Q16)^25^. Members of this family of membrane-integral *b*-type cytochromes typically coordinate two heme *b* cofactors via histidine ligands to the iron ions, giving rise to a proximal heme *b*_P_ and a distal heme *b*_D_ when viewed from the cytoplasmic side. While heme *b*_P_ was present at the expected binding site in *Sa*EMO, the position corresponding to heme *b*_D_ was not occupied by only a single heme group, but by a tightly stacked pair of porphyrin rings arranged in parallel that were separated by only the minimal sterically permissible distance (Fig. 2c,d, Supplementary Fig. 3a). We designate this unprecedented, stacked heme *b* cofactor assembly the twin-heme cofactor, consisting of the stacked heme *b*_D1_ and heme *b*_D2_. Although structurally similar to *Sa*EMO, *Ms*EMO (PDB 9OJN)^21^ also deviates from the canonical arrangement of NarI-like TM domains, in this case by the complete absence of heme *b*_D_. Similarly, the periplasm-facing cytochrome *b* subunit of *Escherichia coli* hydrogenase 1 (PDB 4GD3)^26^ lacks heme *b*_P_. This variability likely reflects the primary function of the TM domains of *Ms*EMO and hydrogenase 1, which is to mediate electron transfer to the MK pool from opposite sides of the membrane without direct coupling to the proton gradient.

**Figure 2.**
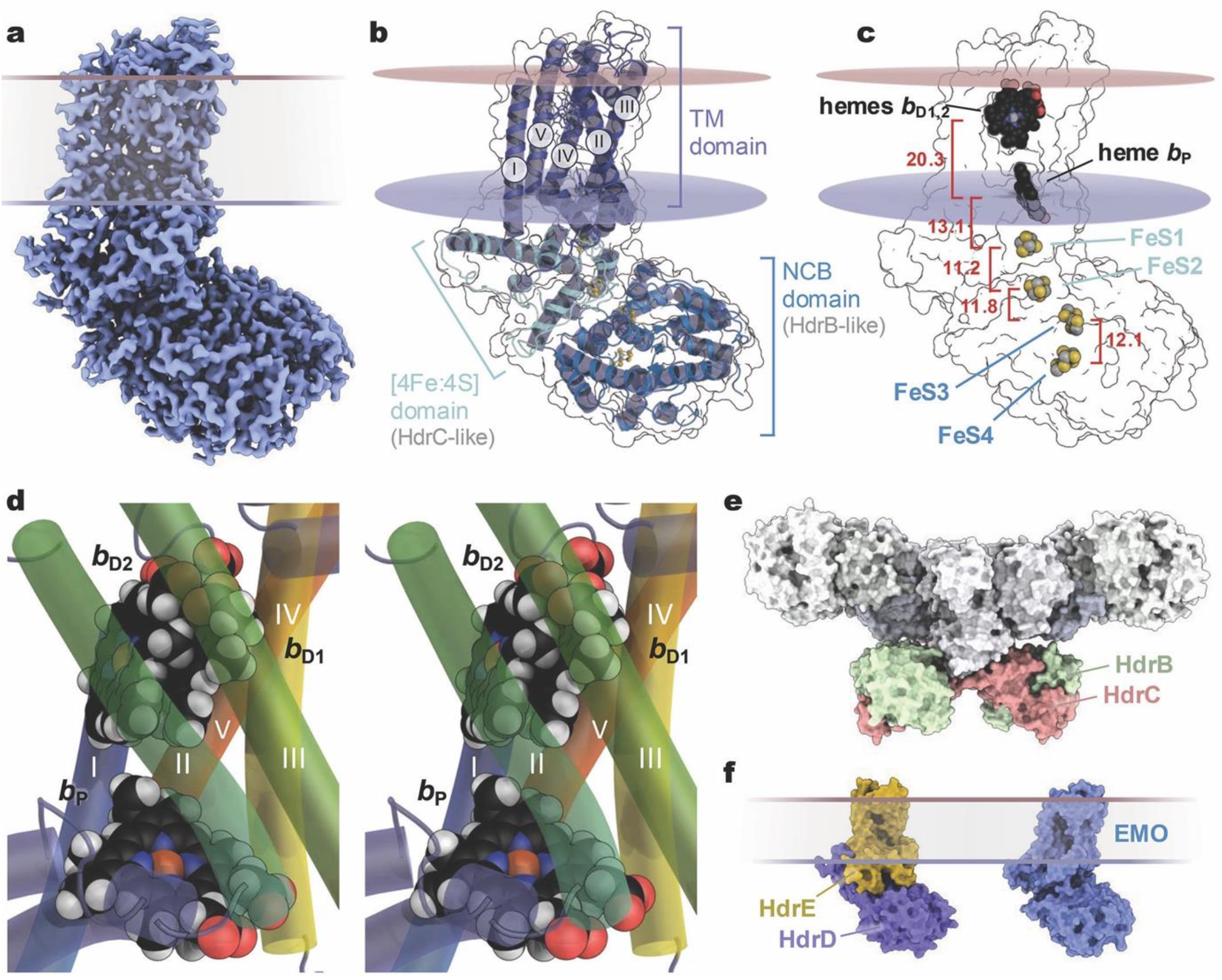
Three-dimensional structure of *Sa*EMO. **a**, Coulomb map at 2.0 Å resolution for *Sa*EMO and its orientation in the cytoplasmic membrane. **b**, Cartoon representation of *Sa*EMO. The single chain is arranged in three distinct domains. The transmembrane (TM) helices I-V are numbered. **c**, Cofactors of *Sa*EMO as denoted in the text. Inter-cofactor distances are given in Ångström (red). **d**, Stereo representation of the three heme cofactors in the TM domain of *Sa*EMO. The TM helices I-V are numbered. **e**, Structure of the soluble HdrABC heterodisulfide reductase (PDB 5ODR) with the prominent HdrB (green) and HdrC (red) subunits. **f**, AlphaFold3 model of the membrane-bound heterodisulfide reductase HdrDE from *M. acetivorans* (left), compared to *Sa*EMO (right).

The two cytoplasmic domains of EMO are homologous in sequence and structure to the HdrB and HdrC subunits of the soluble heterodisulfide reductase complex HdrABC from hydrogenotrophic class I methanogens (Fig. 2e). The α-helical [4Fe:4S] cluster domain comprising residues 261–405, homologous to HdrC, coordinates two canonical [4Fe:4S] cubane clusters (FeS1 and FeS2). This domain connects via a linker (residues 406–450) to the non-cubane cluster-binding (NCB) domain, formerly designated as a ‘cysteine-rich CCG domain’^27^ (residues 451–706), related to HdrB (Fig. 2b,c,e). In HdrABC, the HdrB subunit harbours the active site for the reductive cleavage of the coenzyme M (CoM)–coenzyme B (CoB) heterodisulfide generated by methyl-CoM reductase during methane formation. Structural studies of soluble HDR^23,28^ revealed binding sites for CoM/CoB at the two iron-sulfur clusters of the HdrB subunit, as the first examples of a non-cubane [4Fe:4S] cluster architecture^23^. These clusters, FeS3 and FeS4, are retained in both *Sa*EMO and *Ms*EMO. However, the distal cluster FeS4 displays a remarkable degree of variability. In *Ms*EMO, it forms a [3Fe:4S] cluster, whereas in *Sa*EMO the FeS4 site lacks one cysteine ligand but incorporates an additional sulfide, resulting in a [4Fe:5S] cluster. The FeS1-4 and heme *b*_P_/heme *b*_D1,2_ cofactors are positioned at distances compatible with electron transfer at physiological rates (Fig. 2c). Although the Fe atoms of heme *b*_P_ and heme *b*_D1,2_ are at approximately 20 Å distance, the edge-to-edge distances of the porphyrin systems are considerably shorter.

Full-length *Sa*EMO is predicted to be most similar in size and sequence to membrane-bound HdrDE from class II methanogens. An *in silico* model of *Methanosarcina acetivorans* HdrDE was generated using AlphaFold3^29^ (*Ma*HdrDE) (Fig. 2f), and it was used for comparison to *Sa*EMO (Supplementary Fig. 4a). Notably, the overall structure of *Ma*HdrDE closely resembles that of EMO, with the HdrD subunit forming a fusion of the HdrBC module into a single polypeptide chain, as observed in *Sa*EMO (Fig. 2f). HdrE is an integral membrane protein predicted to contain five membrane-spanning helices. Their arrangement closely resembles that of *Sa*EMO, but differs from *Ms*EMO (see below).

### A distal twin-heme cofactor as a low-potential one-electron carrier for MMK reduction

Positioned analogously to the single heme *b* of *Ms*EMO, the proximal heme *b*_P_ is axially coordinated by residues His87 and H251 in *Sa*EMO (Fig. 3a, Supplementary Fig. 3b). In the AlphaFold3 model of *Ma*HdrDE, it is similarly coordinated by H111 and H249 of HdrE. At the distal side of the membrane, *Sa*EMO contains the stacked twin-heme cofactor, with heme *b*_D1_ coordinated by a single axial histidine, H233 (Fig. 3b). In a multiple sequence alignment, all three histidine ligands are conserved across EMO-like TM domains (Supplementary Fig. 5), including *Ms*EMO, where H223 is present despite the absence of a distal heme in the cryo-EM structure^21^. The two EMO TM domains are topologically equivalent, comprising five TM helices and two short connecting helices after TM helices I and IV. However, TM helix II deviates significantly: while their N-termini align, the C-termini differ by >10 Å. This part of the structure also shows increased flexibility in *Sa*EMO, as highlighted by increased temperature factors of the refined structural model (Fig. 3c). In *Ms*EMO, TM helix II occupies the position of twin-heme *b*_D2_, preventing H223 from coordinating heme *b*_D1_ (Supplementary Fig. 4b). The AlphaFold model of *Ma*HdrDE resembles *Sa*EMO, and the program readily placed three heme groups into the TM domain, one at the *b*_P_ position, and two at the distal site, exactly reproducing the twin-heme arrangement of *Sa*EMO (Supplementary Fig. 4a,c). Remarkably, no distal heme group was placed in an MsEMO model by AlphaFold3, and TM helix II was placed exactly as seen in the experimental structure.

**Figure 3.**
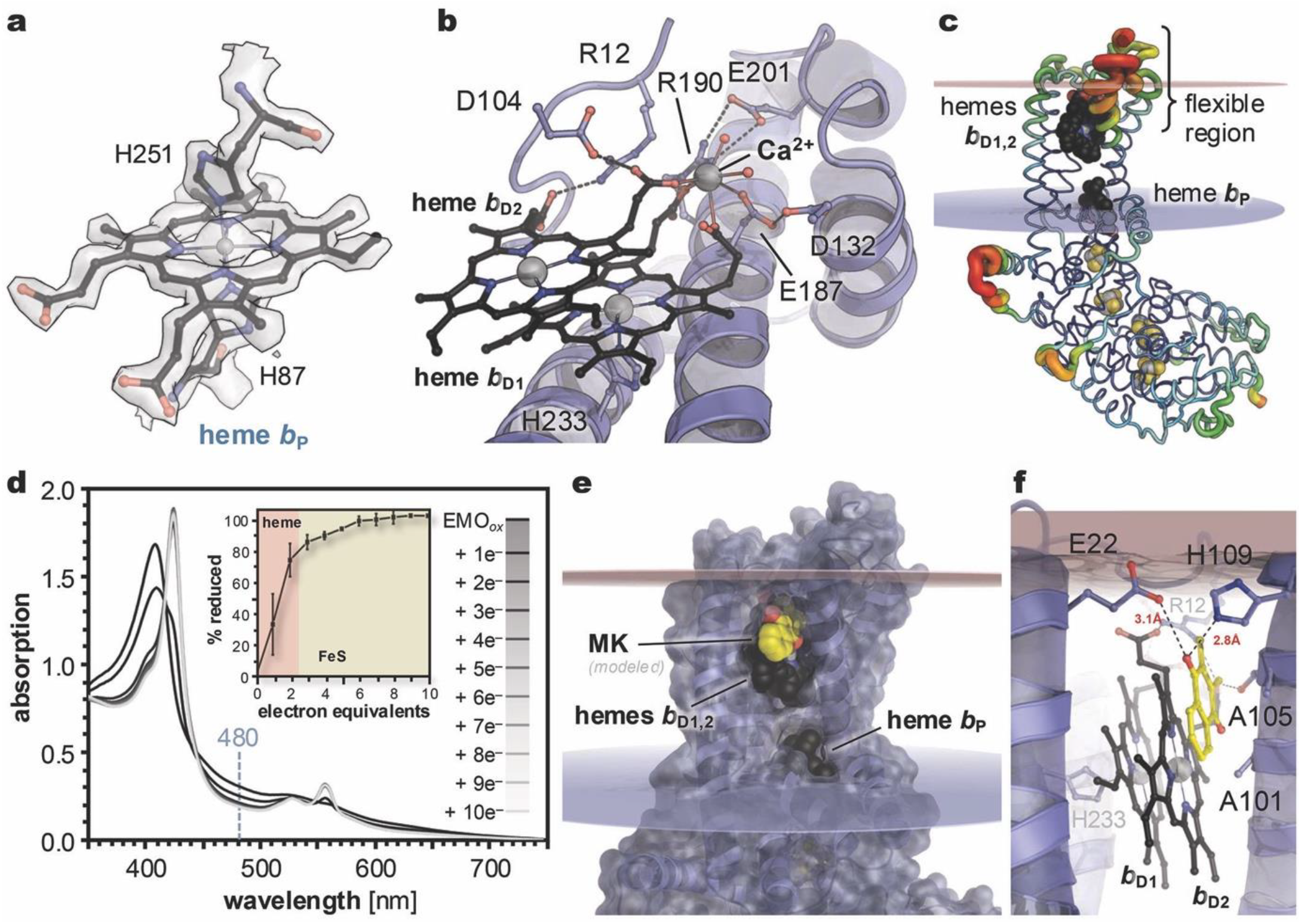
Structural and UV/vis spectroscopic analysis of heme groups, MMK binding and proton transfer during the reverse redox loop of *Sa*EMO. **a**, Heme *b*_P_,coordinated axially by H87 and H251, which preclude MMK binding. **b**, The stacked heme *b*_D1_ and heme *b*_D2_ groups with their coordination environment. **c**, B-factor colouring for *Sa*EMO. Regions of increased flexibility include external loops in the cytoplasmic domain and prominently also the periplasmic region between TM helices II and III that cover the twin-heme cofactor. **d**, UV/vis absorption spectra of *Sa*EMO (8 µM) during dithionite titration. The inset shows reduction followed at 480 nm. Full heme *b* reduction requires two electron equivalents, while complete FeS cluster reduction requires an additional four electrons. **e**, The modelled MMK binding pocket in proximity to heme *b*_D2_. **f**, Docking model for the dimethylnapthoquinone core of MMK at the distal heme *b*_D1,2_. Residues E22 and H109 are in a suitable distance to act as proton donors to the quinone.

The close stacking of the distal twin-heme group in *Sa*EMO and *Ma*HdrDE is highly unusual. The porphyrin rings are arranged in parallel with an interplanar spacing of ~3.6 Å, comparable to the base stacking distance in B-form DNA. Parallel tetrapyrrole stacking is known from multiheme *c*-type cytochromes^30,31^ and the special-pair chlorophylls of photosynthetic reaction centers^32^, but in *Sa*EMO even the iron atoms are separated only by 5.1 Å, making the presence of a bridging atom highly unlikely (Fig. 2d, 3b, Supplementary Fig 3a). Three of the four propionate groups additionally coordinate a well-defined cation, modelled as Ca^2+^, whose octahedral coordination sphere is completed by residue E187 and two water molecules (Fig. 3b). The twin hemes are wedged below TM helices II and III that in the cryo-EM structure exhibited increased flexibility (Fig. 3c). This very close proximity of two hemes suggests electronic coupling, and in line with this, previous redox titrations detected only two single-electron redox transitions at *E*^0^’ = –80 mV for the proximal heme *b*_P_ and one at *E*^0^’ = –220 mV assigned to the distal heme *b*_D_^14^. These observations imply that the twin-heme functions as a single one-electron redox cofactor with an unusually low potential. Dithionite titration of *Sa*EMO confirmed this interpretation: the heme *b* groups were fully reduced after addition of two electron equivalents, whereas further additions caused only minor spectral changes attributable to [4Fe:4S] cluster reduction (Fig. 3d). In total six electron equivalents yielded >95% reduction, consistent with the reduction of four FeS clusters and two heme *b* cofactors.

The high-quality electron density map of *Sa*EMO suggests an amino-acid-derived ligand at the distal axial position of heme *b*_D2_ (Supplementary Fig. 3a). However, the adjacent residue is A101, although the density would be consistent with a serine coordinating the heme *b*_D2_ via its Oγ atom. LC-MS/MS analyses following tryptic and chymotryptic digestion of *Sa*EMO showed no evidence of post-translational modification at this position (Supplementary Fig. 6). Interestingly, the place of A101 in *Sa*EMO is occupied by a serine in the *Ma*HdrDE model that was indeed predicted to act as a distal ligand to heme *b*_D2_ (Supplementary Fig. 4c), rendering the observed density in the *Sa*EMO structure even more enigmatic.

### Potential MMK binding and proton transfer during reverse electron transfer

In the proposed reverse redox loop of *Sa*EMO, MMK is reduced near the distal twin-heme to harness the proton gradient (Fig. 1), and extraction from purified *Sa*EMO followed by ultra-performance liquid chromatography analyses indeed yielded 0.6–0.7 mol MMK bound per mol *Sa*EMO (Supplementary Fig. 7). However, cryo-EM analyses with various MMK derivatives did not resolve the binding site, possibly due to the high detergent concentrations needed to solubilize EMO from the membrane. Structural characterization of quinone-dependent oxidoreductases with native quinones remains generally challenging, as most structures have been obtained with inhibitors, often leaving details of the physiological quinone site uncertain.

For the *Ms*EMO structure, the authors modelled a bound MK next to heme *b*_P_, in the center of the membrane^21^. This position is unusual for a quinone interacting with redox cofactors, as it does not provide a proton pathway to either side of the membrane. The corresponding pocket exists in *Sa*EMO but is occluded by the repositioned F94 and F240. In *Sa*EMO, MMK likely binds in proximity to hemes *b*_D1,2_ and a pocket above heme *b*_D2_ is sufficiently large to accommodate the dimethyl naphtoquinone moiety with its methyl groups orientated inward (Fig. 3e). Molecular modelling suggests that a slight displacement of TM helix II permits MMK binding, which likely remains shallow, consistent with the lack of density at high detergent concentrations. On the periplasmic side, the pocket is lined by residues E22 and H109 which may function as proton conduits for MMK reduction (Fig. 3f). In the modelled state, one MMK oxygen can be protonated via H109 or E22, both of which access the periplasm, while, the second oxygen of the *para*-quinone is positioned near the central iron ion of heme *b*_D2_ (Fig. 3f).

### The [4Fe:5S] and [4Fe:4S] non-cubane clusters exhibit unusual spectroscopic properties

In HDR, the non-cubane [4Fe:4S] clusters HB1 and HB2 are typically coordinated by 10 cysteine residues within the NCB domain (Fig. 4a). Crystal structures of HdrABC revealed that these FeS clusters constitute the catalytic site for heterodisulfide reduction, each binding either the CoM or CoB substrates (Fig. 4a)^23^. Although the cytoplasmic domain of *Sa*EMO closely resembles HdrBC (Fig. 2b,c) and HdrD (Fig. 2f), it lacks the open substrate-binding cavity of HDR. In *Sa*EMO, H534 occupies the free CoM-binding site of the proximal FeS3 (Fig. 4b). In contrast, the distal non-cubane FeS cluster FeS4 differs by replacement of a cysteine ligand (C81 in HDR from *M. thermolithotrophicus*) with G535, such that an inorganic sulfide occupies the position corresponding to the HdrB cysteine thiolate (Fig. 4b), resulting in an unprecedented non-cubane [4Fe:5S] cluster. The origin of the additional fifth sulfide remains unclear. In *Ms*EMO, FeS3 corresponds to a canonical non-cubane [4Fe:4S] cluster, whereas substitution of C42 in *M. thermolithotrophicus* HdrB by T539 gives rise to a linear non-cubane [3Fe:4S] cluster (Fig. 4c)^21^.

**Figure 4.**
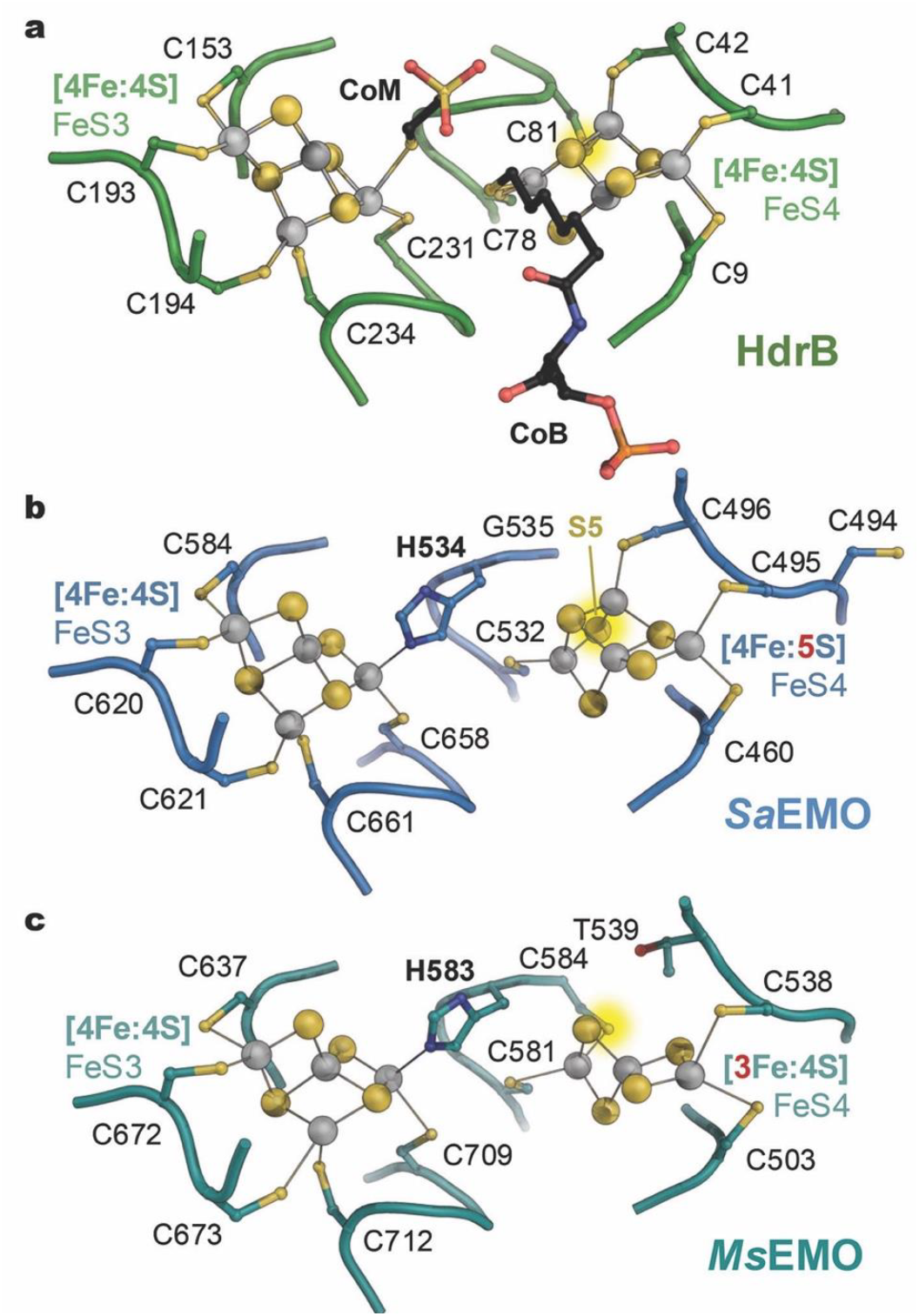
Structural variability in non-cubane iron-sulfur clusters in NCB domains. **a**, Active site of *M. thermolithotrophicus* heterodisulfide reductase (PDB 5ODR), with the bound coenzymes B and M and the two non-cubane [4Fe:4S] clusters. **b**, The *Sa*EMO clusters FeS3 and FeS4. H534 blocks the position that serves as a substrate binding site in HDR, while the distal FeS4 cluster has expanded into a [4Fe:5S] unit, compensating for an absence of C535. **c**, *Ms*EMO (PDB 9OJN) features a [4Fe:4S] FeS3 cluster with a H583 ligand. In FeS4, C539 is substituted with threonine, resulting in loss of an iron ion and the formation of a linear [3Fe:4S] cluster.

To further analyse the non-cubane clusters of *Sa*EMO, we heterologously produced the NCB domain of the protein (E406-A706, Fig. 5a), yielding a highly enriched, soluble 36 kDa protein (Fig. 5b). Iron quantification (7.8 ± 1.3 mol Fe per protein) aligns with the [4Fe:4S] FeS3 and [4Fe:5S] FeS4 non-cubane clusters identified in the cryo-EM structure of *Sa*EMO. The UV/vis absorption spectrum was more reminiscent of [2Fe:2S] clusters, showing a pronounced shoulder at 550 nm and a smaller feature at 720 nm (Fig. 5c). Reduction with dithionite bleached the spectrum after the addition of two electron equivalents (Fig. 5c), indicating that both non-cubane clusters function as single-electron redox cofactors.

**Figure 5.**
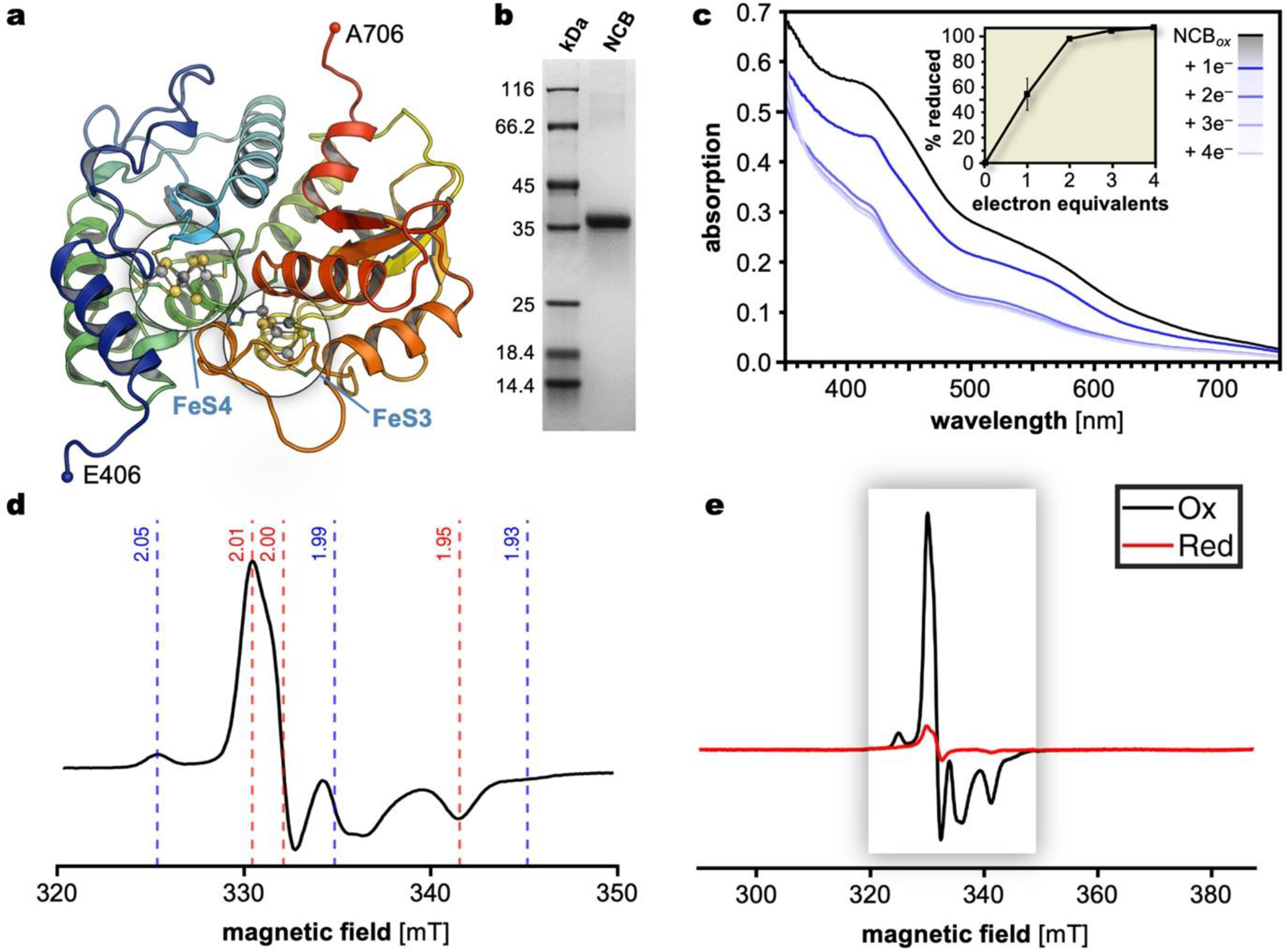
Spectroscopic properties of heme *b* cofactors of EMO and the non-cubane clusters of heterologously produced NCB domain. **a**, Structural model of the isolated NCB domain of SaEMO that was used for spectroscopic characterization (partial structure from *Sa*EMO). **b**, SDS-PAGE of the isolated, 36 kDa NCB domain after two chromatographic steps. **c**, UV/vis spectroscopic titration of the NCB domain with dithionite to monitor cofactor reduction. The inset shows the extent of reduction quantified from the decrease in absorbance at 420 nm. **d**, EPR spectra of the NCB domain at 40 K and 0.63 mW in the oxidized (as isolated) state. The principal *g*-values of the two spin species are shown in red for the species 1 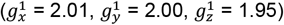 and in blue for the species 2 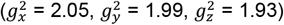. **e**, EPR spectra at 40 K and 0.63 mW in the oxidized (black) and the dithionite-reduced state (red).

The non-cubane clusters of the NCB domain were further characterized by electron paramagnetic resonance (EPR) spectroscopy. In the as-isolated, oxidized state, prominent rhombic signals were observed in the *g* = 2 region up to 50 K that are rather characteristic of [2Fe:2S]^+^ clusters. Varying the temperature and microwave power resolved two distinct *S* = 1/2 species and their principal *g*-values (Supplementary Fig. 8). Unlike cubane [4Fe:4S] clusters, both clusters show a fast power saturation at lower temperatures (10 K or 20 K, Supplementary Fig. 8, 9). Species 1 showed an almost axially symmetric *g*-tensor 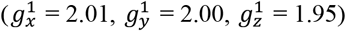, while species 2 displayed a broader, ortho-rhombic *g*-tensor (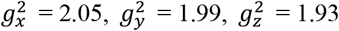;Fig. 5d). Oxidation with dichlorophenol indophenol did not alter the spectra, indicating the clusters were in the paramagnetic [4Fe:4S]^3+^ and [4Fe:5S]^3+^ states. Dithionite reduction strongly diminished the EPR signals, consistent with the formation of diamagnetic [4Fe:4S]^2+^ and [4Fe:5S]^2+^ species (Fig. 5e), and allowed for a distinction between the two species, since only the species 2 signal was completely abolished (Supplementary Fig. 10). Although a definitive assignment of the two EPR species to the [4Fe:4S] and [4Fe:5S] non-cubane clusters was not yet possible, the UV/vis and EPR spectra consistently indicate features reminiscent of [2Fe:2S] clusters cycling between the +3 and +2 redox states. Both the UV/vis and EPR spectra are similar to the non-cubane [4Fe:4S] clusters from *Desulfovibrio vulgaris* Hildenborough HdrB, which showed *g*_xyz_-values of 2.02, 1.99, and 1.95, although the iron occupancy was only ~50% ^33^. In contrast, HDR from *Methanothermobacter marburgensis* exhibited a similar signal with *g*_xyz_-values of 2.01, 1.99, and 1.94 only in the presence of CoM^34^.

### Conformational flexibility in the EMO:ETF complex ensures efficient electron transfer from FAD to the non-cubane clusters

To assess the formation of a complex between *Sa*EMO and its interaction partner *Sa*EtfAB (Fig. 1), the enriched proteins were co-incubated (1:1) and analysed by Blue Native (BN) PAGE. A novel protein band corresponding to a molecular mass of ~200 kDa appeared only upon co-incubation, as visualized by Coomassie and in-gel heme staining (Supplementary Fig. 11). LC–MS/MS identified EtfA (57.8%), EtfB (53.3%), and EMO (24.7%). The predicted 1:1:1 complex (~160 kDa) appeared larger in BN-PAGE due to the detergent micelle of *Sa*EMO. This suggested complex formation likely facilitates electron transfer from FAD in *Sa*EtfA to the non-cubane FeS clusters in *Sa*EMO. The complex presumably is transient, as *Sa*ETF also engages acyl-CoA DH partners. Consistent with this assumption, no ternary EMO:EtfAB:cyclohexanoyl-CoA DH (a typical acyl-CoA DH from *S. aciditrophicus*^24^) complex was observed.

We then proceeded to determine a single-particle cryo-EM structure of the *Sa*EMO:EtfAB complex to 3.0 Å resolution (Fig. 6a, Supplementary Table 1, Supplementary Fig. 12). Here, *Sa*EtfAB binds to the cytoplasmic domain of *Sa*EMO exclusively via the EtfB subunit. *Sa*EtfAB binds an AMP nucleotide in subunit B that likely represents a vestige of the FAD cofactor found in electron-bifurcating ETFs^35^. Consistent with known ETF structures, EtfA comprises an N-terminal βαβ-domain and a C-terminal Rossman-fold domain that binds FAD. In free ETFs, the C-terminal domain of EtfA positions its FAD moiety toward the AMP ligand of EtfB, thereby shielding the cofactor and preventing both dissociation and electron transfer. Within the *Sa*EMO:ETF complex, however, the FAD domain of EtfA was rotated by approximately 60° towards the surface of *Sa*EMO (Fig. 6b), positioning it at a distance suitable for electron transfer (Supplementary Movie 1). In the cryo-EM reconstruction, the 7- and 8-methyl groups of the isoalloxazine ring of FAD are nearly equidistant (12 Å *vs*. 13 Å) to the clusters FeS3 and FeS4, suggesting that either one could serve as the initial electron acceptor. The reduced local resolution of the FAD reflects conformational flexibility, and we cannot exclude the possibility that FAD transiently approaches one of the terminal clusters of EMO even more closely to adopt an optimal configuration for electron transfer (Fig. 6c).

**Figure 6.**
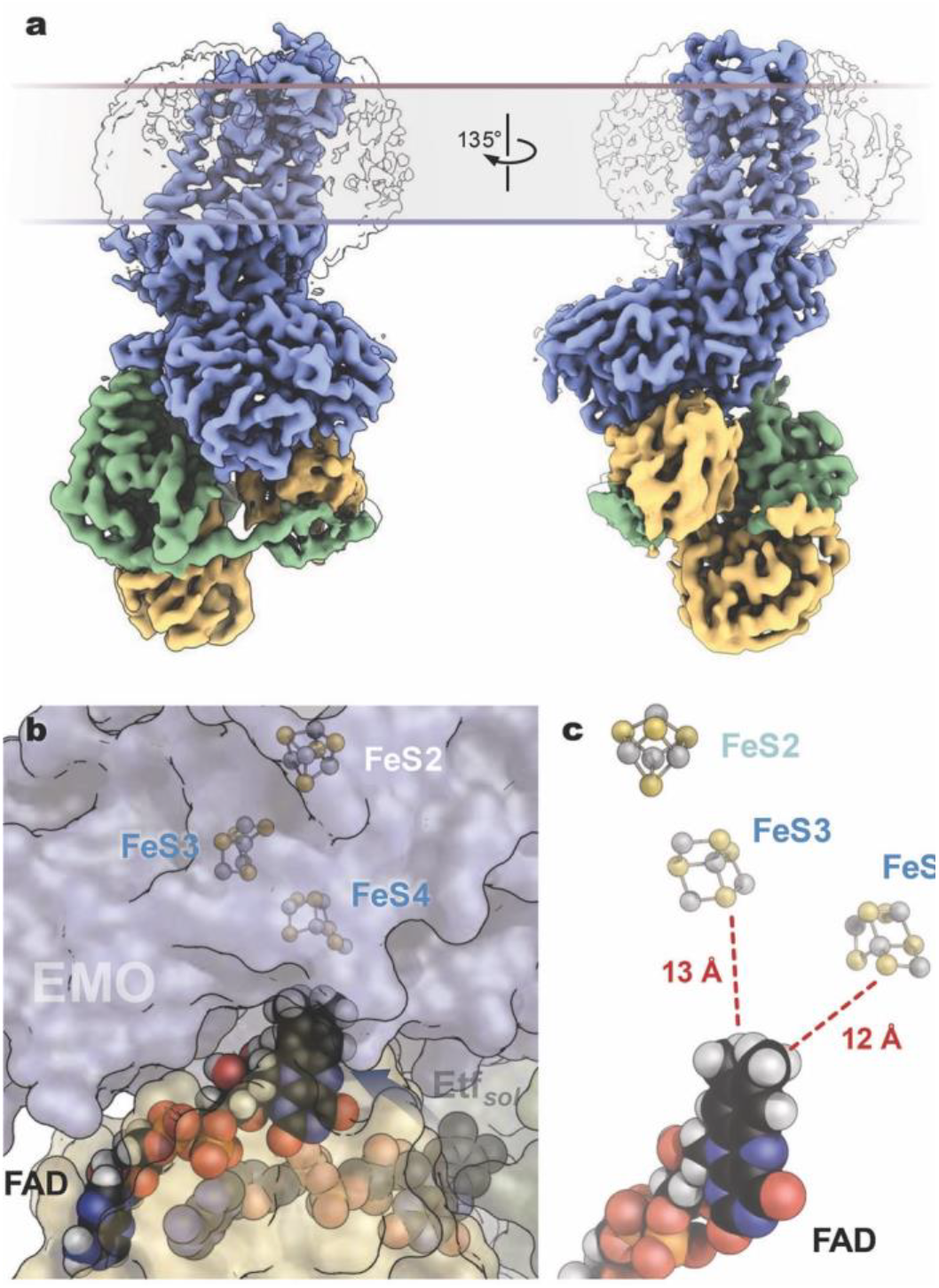
Structure of the EMO-ETF complex. **a**, Coulomb map for the 3.0 Å resolution model of the *Sa*EMO:EtfAB complex in two orientations. **b**, The FAD cofactor of EtfAB and its positioning relative to the distal iron-sulfur clusters of *Sa*EMO. The transparent FAD moiety denotes its position in an AlphaFold model of the closed conformation of soluble ETF, indicating the 60° rotation that the FAD domain must undergo to initiate electron transfer (Supplementary Movie 1). **c**, In the conformation observed in the experimental structure, the isoalloxazine moiety of FAD is nearly equidistant to clusters FeS3 and FeS4 of EMO.

### An evolutionary link between methanogenesis and fatty acid β-oxidation

The presence of non-cubane clusters in EMO is striking, given their highly specialized role in catalysis in HDR. In EMO, these clusters do not serve as catalytic sites but mediate electron transfer from reduced ETF to the (M)MK pool. Notably, HDR and experimentally characterized EMO from *S. aciditrophicus* and *M. smegmatis* only share the proximal non-cubane cluster (HB1/FeS3, Fig. 4), while the distal cluster shows an intriguing degree of variability (Fig. 4).

To assess the distribution and evolutionary links of NCB domains in methanogenesis, fatty acid β-oxidation and beyond, we conducted a phylogenetic analysis considering the predicted or known functions of their associated modules (Supplementary Table 2, Supplementary Fig. 13, 14). Across these oxidoreductases, NCB domains are consistently connected to a soluble domain with two cubane [4Fe:4S] clusters. EMO and membrane-bound HDR (HdrDE) form distinct phylogenetic clusters, each combining soluble NCB and cubane domains with a tri-, di- or mono-heme *b* membrane domain, reflecting their roles in electron transfer from ETF_red_ to (M)MK or from reduced methanophenazine to CoM-S-S-CoB^36^. In archaeal succinate dehydrogenase, NCB domains may mediate electron transfer from succinate to quinones (or reverse), although no heme *b*-containing TM domain is present^37,38^. In thiol:fumarate reductase, TfrB, CoM/CoB thiols acts as electron donors for fumarate reduction instead of membrane-bound carriers, indicating a catalytic role for the NCB domain^39^. In other NCB-containing enzymes, including soluble HDRs (HdrB)^23,28^ and class II benzoyl-CoA reductases (BamD)^40-42^, NCB and cubane domains are linked to additional electron-transfer subunits, often facilitating electron flow to quinones, as is the case in EMO. The function of NCB domains is less defined in other clades such as GlcF^43^, D-iLDH^44^, and GlpC^17^. The latter is a subunit of anaerobic glycerol-3-phosphate DH in *E. coli*, which may help to rationalize the ability of this bacterium to heterologously produce the *Sa*EMO NCB domain with high cluster occupancy. The absence of auxiliary genes suggests that no dedicated assembly machinery is required for NCB cluster biosynthesis.

## Conclusion

This study provides first structural insights into a central energy-coupling process that links biomass-derived fatty acid oxidation to methane formation. Central to this process is an unusual low-potential twin-heme cofactor that functions as a single-electron transfer unit, enabling proton-motive-force-driven electron transfer from reduced ETF to methylmenaquinone. The unique redox properties of this cofactor identify it as a specialized key component for coupling fatty acid oxidation to methanogenesis through a reverse redox loop. Future spectroscopic analyses of targeted EMO variants will be essential to elucidate the molecular basis of twin-heme-mediated energy transduction.

Beyond its significance in biomass-to-biogas conversion, EMO provides insights into an evolutionary connection with the membrane-bound methanogenic heterodisulfide reductase complex HdrDE. Structural conservation of the membrane and non-cubane-cluster domains, together with phylogenetic analyses, suggests that EMO originated from an ancestral methanogenic enzyme that underwent a functional transition from heterodisulfide reduction to ETF- and quinone-linked electron transfer. This evolutionary shift converted a catalytic redox module into a specialized electron-transfer system, underscoring the remarkable adaptability of NCB-containing oxidoreductases. More broadly, these findings establish a mechanistic and evolutionary link between methanogenic energy conservation and fatty acid β-oxidation, illustrating how ancient redox modules were repurposed to support emerging metabolic functions.

## Materials and Methods

### Bacterial strains and plasmids

*S. aciditrophicus* strain SB (DSMZ 26646) was grown on crotonate as sole carbon and energy source at a 200 L scale in anaerobic mineral salt medium (pH 7.3) as described before^24^ and stored in liquid nitrogen. *E. coli* BL21(DE3) Δ*iscR*^45^ was used for the heterologous production of the NCB domains of EMO. The strain was transformed (heat shock) with a pPR-IBA 101 plasmid (IBA-Lifesciences) containing the truncated *emo* gene (SYN_02638, E406-A706) fused to a C-terminal Twin Strep tag II for affinity enrichment under control of a T7 promotor. Transformed *E. coli* BL21(DE3) Δ*iscR* containing a plasmid for the heterologous production of the NCB domains of EMO (E406-A706) was cultivated in LB medium containing 50 mM 3-(*N*-morpholino)propanesulfonic acid (MOPS) at a 2L-scale as described elsewhere for the heterologous production of recombinant HdrB^46^. Protein production was induced with 400 µM isopropyl β-D-thiogalactoside (IPTG) at optical densities of approx. 0.5. Cells obtained after 20 h of anaerobic cultivation at 20 °C were harvested anaerobically by centrifugation and stored at –20 °C until further use. Cyclohexanoyl-CoA DH of *S. aciditrophicus* was heterologously produced in *E. coli* BL21 (DE3) Star OneShot as described previously^24^.

### Protein Isolation

All protein enrichment steps were carried out in an anoxic atmosphere (95% N_2_, 5% H_2_) at 8 °C. Purity analyses throughout enrichment were performed using sodium dodecyl sulphate-polyacrylamide gel electrophoresis (SDS-PAGE) with 12% acrylamide gels.

#### EMO and ETF

For the enrichments of EMO and EtfAB from approximately 10 g of *S. aciditrophicus* wild-type cells, established protocols^14^ were followed. Briefly, enrichment of EMO from *S. aciditrophicus* membranes was achieved by solubilisation with *n*-dodecyl-β-*D*-maltopyranoside (DDM) followed by anion exchange and size-exclusion chromatography. For the enrichment of ETF from the soluble protein fraction, anion exchange chromatography followed by ammonium sulphate precipitation and hydrophobic interaction chromatography were employed. Concentrated protein aliquots were stored in anoxic containers at –70 °C until further use.

#### NCB domain of EMO

Cells were resuspended in a two-fold volume of buffer A (50 mM Tris/HCl, 150 mM NaCl, pH 8.0) in the presence of a spatula tip of DNase I, dithioerythritol (DTE) and lysozyme, each. Disruption was achieved using a French pressure cell (1,100 psi) and followed by the removal of insoluble cell components and debris by ultracentrifugation for 1 h at 150,000 *×g*. Soluble, Strep-tagged NCB domains were enriched by Strep-Tactin XT (IBA-Lifesciences) affinity and subsequent Superdex 75 Increase (Cytiva) size-exclusion chromatography in buffer A. Elution during affinity chromatography was achieved using 50 mM biotin in buffer A. Protein samples were concentrated (10 kDa cut-off) and stored in anoxic containers at –70 °C until further use.

#### Cyclohexanoyl-CoA DH

Enrichment of cyclohexanoyl-CoA DH via histidine tag affinity chromatography was performed following established protocols^23^. Concentrated protein aliquots were stored in anoxic containers at –70 °C until further use.

### Determination of electron transfer from acyl-CoA to EMO

Electron transfer from cylohexanoyl-CoA (synthesized as described elsewhere^47^) to the proximal heme *b*_P_ of EMO via ETF and cyclohexanoyl-CoA DH were detected by UV/vis spectroscopy under anoxic conditions (N_2_/H_2_ atmosphere) *in vitro*, using a UV-1800 UV/vis spectrophotometer (Shimadzu) and quartz cuvettes. Measurements were carried out in buffer A. Spectra were recorded from 200 to 800 nm, baseline-corrected at 800 nm and corrected for dilution effects. The reduction of heme *b* groups was monitored by calculating and plotting the absorption differences between 413 and 427 nm (Soret band) and between 560 and 580 nm over time (30 min in total). The reaction mix contained 3 µM of DDM-solubilized oxidized EMO, 6 µM ETF, 2 µM cyclohexanoyl-CoA DH and 100 µM cyclohexane-CoA.

### Blue Native-PAGE analysis of the EMO–ETF interaction

Complex formation between *Sa*EMO and *Sa*EtfAB was investigated under anoxic conditions and evaluated via BN-PAGE using 4–16% acrylamide gradient gels (SERVA Electrophoresis GmbH) in the presence of 0.003% (*w/v*) of DDM added to the cathode buffer (15 mM bis-Tris, 50 mM tricine, 0.001% (*w/v*) Coomassie G250). The anode buffer consisted of 50 mM bis-Tris (pH 7). Samples were prepared by mixing enriched and concentrated EMO and ETF samples in the desired molar ratio, followed by incubation for 15 min at room temperature. Directly before loading, standard BN-PAGE loading buffer (4-fold stock) was added. Following electrophoresis at 300 V, in-gel heme staining^48^ was carried out before Coomassie staining of proteins. For these purposes, proteins were first fixed using a solution containing 30% (*v/v*) methanol and 20% (*v/v*) acetic acid for 10 min. Excess Coomassie dye from electrophoresis was removed by incubating the gel in a mixture of 30% (*v/v*) of 2-propanol and 70% (*v/v*) 250 mM sodium acetate buffer (pH 5) for 30 min, twice. Next, the gel was incubated in 20 mL of a solution of 30% (*v/v*) tetramethylbenzidine (3 mM) in methanol and 70% (*v/v*) 250 mM sodium acetate buffer (pH 5) for 1 h in the dark. 200 µL H_2_O_2_ (3 M) were added and after another 30 min of incubation in the dark, protein-bound heme groups exhibited a light blue colour. Afterwards, proteins were stained using Coomassie (FastGene Q-stain, Nippon Genetics). For identification of protein bands putatively containing interacting proteins by mass spectrometry (MS), the respective protein bands were excised, reduced, alkylated and treated with trypsin. The resulting peptides were then separated and analysed via an I-class ultra-performance liquid chromatography (UPLC) system (Waters) coupled to a Waters Peptide Synapt G2-Si HDMS ESI/Q-TOF mass spectrometer and matched with the peptide sequences of *S. aciditrophicus* strain SB, as described^49^.

### Cryo-EM data collection, processing and model building

Datasets for both *Sa*EMO and the *Sa*EMO:ETF complex were collected. The EMO sample was diluted to 10 mg mL^−1^ prior to preparing the grids. For the EMO:ETF complex, the purified samples of EMO and ETF were mixed in a 1:1 molar ratio in an anaerobic glove box, then the mixture was split into three fractions: one aliquot was reduced with 5 mM sodium dithionite, one was oxidized with 1 mM potassium ferricyanide, and the third one was untreated. Cryo-EM grids were prepared using a Vitrobot Mark IV (Thermo Fisher Scientific) as follows: 3 µL of the protein sample was applied to glow-discharged Quantifoil Cu R1.2/1.3 300-mesh grids, waited for 5 s and blotted for 6 s with filter paper, and plunge-frozen in liquid ethane cooled by liquid nitrogen. Data collection was performed on a 300 kV Krios G4 cryo-TEM (Thermo Fisher Scientific) equipped with a Falcon 4i direct electron detector. Images were acquired using a 10 eV slit width on the energy filter, with a total dose of 40 e^−^/Å^2^ in electron-event representation (EER) format.

The micrographs were processed in cryoSPARC v4.7 (Supplementary Fig. 2, 12)^50^. The EER data were imported (40 fractions), motion corrected and contrast transfer function (CTF) estimated. Initial particle picking employed a blob picker, followed by extraction with down-sampling and 2D classification. Selected particle subsets underwent *ab-initio* reconstruction to generate reference volumes for heterogeneous refinement. Concurrently, representative 2D classes were used for template-based picking, with subsequent particles processed through two rounds of heterogeneous refinement. The particles of the best 3D classes were combined and re-extracted without down-sampling. Additional rounds of hetero-refinement and 3D classifications were performed to further remove bad particles, and the best class was subjected to non-uniform refinement^51^. The EMO dataset was processed to 2.0 Å resolution (Supplementary Fig. 2). The datasets of the EMO:ETF complex resulted in maps at 3.0 Å resolution. The AlphaFold model of the EMO:ETF complex was fitted into the

3.0 Å map. The density for the FAD-binding domain of ETF was relatively weak, indicating its dynamic nature. Thus, 3D classification and 3D variability analysis with focused mask were performed to track the multiple conformations of the FAD-binding domain (Supplementary Fig. 12).

The AlphaFold^29^ models of EMO and EMO-ETF were used as starting models for building. The model was fitted into the density map using UCSF ChimeraX^52^, followed by iterative refinement in COOT^53^ and refined in real-space with PHENIX^54^. Structure validation was performed using MolProbity^55^. Data collection and refinement statistics are summarized in Supplementary Tab. 1. Figures were generated using PyMOL (Schrödinger LLC) or UCSF ChimeraX^52^.

### UV/vis spectroscopy of EMO and NCB domain of EMO

Measurements were carried out in buffer A using quartz cuvettes and a UV-1800 UV/vis spectrophotometer (Shimadzu) under anoxic conditions. Spectra were recorded from 300 to 800 nm, baseline-corrected for 800 nm and corrected for dilution effects. EMO was used at 8 µM while the NCB domain of EMO (E406-A706) was used at 20 µM. The proteins were stoichiometrically reduced with sodium dithionite in 1.0 electron equivalent steps, respectively. For this purpose, the reducing capacity of sodium dithionite was determined experimentally by titration of 10 mM methyl viologen with dithionite and recording spectra between 300 and 800 nm. The changes in absorption at 730 nm were used to calculate the actual concentration of reducing equivalents of dithionite added using the molar extinction coefficient of methyl viologen at 730 nm (ε_730 nm_ = 2,400 cm^−1^·;M^−1^)^41^. The concentrations of dithionite needed for the stoichiometric reduction of EMO and the recombinant NCB domains were adjusted accordingly.

### Quinone extraction from enriched EMO samples and UPLC analysis

Quinone species co-enriched with EMO were extracted and analysed by UPLC (Waters H-Class UPLC system). To 5 µL of enriched EMO (8.9 mg·mL^−1^, triplicates), 40 µL ethanol were added and the mix was incubated for 2 h at 30 °C. Denatured protein was removed by centrifugation (14,000 rpm, 20 min). 10 µL methanol was added to the supernatant and 10 µL of the resulting solution were applied to a Waters Acquity CSH-C18 column (2.1 *×*100 mm, 1.7 µm particle size) in triplicates. Separation was achieved at a flow rate of 0.15 mL·min^−1^ at 50 °C column temperature in an isocratic ethanol:methanol (4:1) gradient (6 min). Detection of eluting quinone species was achieved using a photo diode array detector. For quantification, authentic standards of MK-7 (Sigma-Aldrich) at known concentrations (2 to 50 µM) were prepared and analysed using the same procedure (three injections, each) to construct a calibration line (peak areas at 248 nm vs. concentrations). Following this procedure, only peaks exhibiting UV spectra characteristic for (M)MK species obtained from extracted EMO samples were considered for quantification.

### High-resolution mass spectrometric analysis

Purified EMO was denatured in 1% SDS, cysteine residues were reduced with 10 mM DTT for 15 min at 95 °C and alkylated using 50 mM chloroacetamide for 30 min at 22 °C. The reaction was quenched with 50 mM DTT. Digestion with trypsin was performed by a custom SP3 bead-based purification protocol^56^ on a Hamilton STARlet automated liquid handling system (Hamilton). Proteins were incubated with 100 µg of Sera-Mag SpeedBead magnetic carboxylate-modified particles (Cytiva) in 80% ethanol for 20 minutes at room temperature in a 96-deep-well plate (Storage plate 96 well, 1 mL, Agilent Technologies) and separation was conducted using a 96-well magnet plate (Magnum Flex, Alpaqua). After binding, samples were washed three times with 80% ethanol. Proteins were digested overnight at 37 °C in 50 µL of 100 mM ammonium bicarbonate containing 0.15 µg of trypsin (Promega V5111). Digestion was stopped the following day by adding 30 µL of 5% formic acid (FA). For digestion with chymotrypsin, an aliquot of the sample was diluted with 100 mM ammonium bicarbonate to SDS concentration of 0.2%. Chymotrypsin was added in a ratio of 1:25 and digestion preformed at 25 °C overnight. Peptide solutions were desalted using self-packed STAGE tips^57^ composed of three layers of 1.0×1.0 mm SDB-RPS (AttractSPE Disks Bio RPS, Affinisep) equilibrated sequentially with 100% methanol, 80% acetonitrile containing 0.1% FA, and twice with 0.1% FA. Acidified peptide solutions were loaded, washed with 0.1% FA, 80% acetonitrile with 0.1% FA, and 80% methanol with 0.5% FA, followed by elution using 60% acetonitrile containing 5% ammonia. The eluates were dried in a vacuum concentrator and reconstituted in 0.1% FA prior to loading onto Evotips (EV2013 Evotip Pure, Evosep), following the manufacturer’s protocol.

For LC-MS analysis, an Evosep One nano-LC system (Evosep) was coupled online to an Exploris 480 mass spectrometer (Thermo Fisher Scientific). Peptides were separated using a 44-minute gradient (30 SPD workflow) with a 15 cm×150 µm performance column (EV1137, Evosep) maintained at 40 °C using a column oven (PRSVO-V2, Sonation). Electrospray ionization was performed using a Nanospray Flex ion source (Thermo Fisher Scientific) with a stainless-steel emitter featuring an integrated liquid junction (EV1072, Evosep) and mounted into an EasySpray adapter (EV1072, Evosep). A spray voltage of +1800 V was applied, and the ion transfer tube was set to 275 °C. Data were acquired in both data-dependent acquisition (DDA) and data-independent acquisition (DIA) mode. Cycles of DDA consisted of: one MS1 survey scan (RF lens of 40%, normalized AGC target of 300%, maximum injection time of 25 ms, *m/z* range of 350 to 1.400, resolution of 120.000 at 200 *m/z*, profile mode) followed by MS/MS scans on the 12 most abundant peptide signals of the survey scan by higher-energy collision-induced dissociation (HCD) at a normalized energy of 28%, combining an AGC target value of 200% and an MIT of 25 ms, with a mass isolation window of 1.3 *m/z*, a minimum signal intensity threshold of 2×10E5 at a resolution of 15,000, RF level of 50%, exclusion time 45 s and in centroid mode. For acquisition cycles of DIA, the MS1 survey scan had a maximum injection time of 45 ms and was followed by MS2 fragment spectra (RF lens 50%, normalized AGC target 1000%, maximum injection time 54 ms, resolution 30,000, profile mode) generated by HCD with variable isolation widths of 14 *m/z* in the precursor range from 361 to 450 *m/z*, 7 *m/z* in the range from 450 to 800 *m/z* and 14 *m/z* in the range from 800 to 1100 *m/z*. Isolation windows overlapped by 1 *m/z*.

For peptide identification, DDA raw data were matched against the sequence of EMO from *S. aciditrophicus* (SYN_02638) supplemented with sequences of common contaminants using MaxQuant (version 2.0.2.0)^58^. Trypsin/P or Chymotrypsin+ was specified as proteolytic enzyme in semispecific mode with dependent peptides search activated. Oxidation of methionine was set as variable and carbamidomethylation of cysteine as fixed modification. DIA spectra were searched using FragPipe (version 22)^59^ with the integrated MSFragger (version 4.3) search engine considering semispecific peptides generated by Trypsin with a length between 7 and 56, or generated by Chymotrypsin with a length between 5 and 50 allowing the same modifications as for DDA searches. Peptide identifications were filtered at 1% FDR.

### EPR spectroscopy of NCB domain

The heterologously produced NCB domain of EMO (E406-A706) was analysed using EPR spectroscopy. Three samples (114 µM enzyme in buffer A) were prepared: as purified, oxidized by 2,6-dichlorophenolindophenol (DCPIP, 180 µM) and reduced by sodium dithionite (450 µM). Samples were prepared anaerobically and, for the reduced and oxidized samples, incubated at room temperature for 10 min with the reducing/oxidizing agent before flash freezing in liquid nitrogen.

All samples were kept frozen and inserted at 80 K into the EPR cavity. All data sets were collected by using a Bruker ELEXSYS E500 (with digital upgrade) together with a Bruker SuperX bridge and a Bruker SHQ4112 cavity. The sample was immerged in an Oxford E900 cryostat, and the temperature was controlled by an Oxford Mercury ITC. All samples were measured with the following parameters: modulation frequency 100 kHz, modulation amplitude 0.3 mT and a conversion time of 327 ms. The temperature was changed between 10 K and 50 K in 10 K steps. The microwave power was swept over the following values: 20 mW, 6.3 mW, 2.00 mW, 0.63 mW, 0.20 mW, 0.06 mW and 0.02mW. The magnetic field was calibrated by using a DppH (2,2-Diphenyl-1-Picrylhydrazyl) with a known *g*-factor of 2.0036. The cavity and cryostat background was always measured under the same conditions by using a tube with sample buffer. The background was afterwards subtracted. A polynomial baseline correction was also applied to each data set.

### Multiple sequence alignments and phylogenetic analysis

Multiple sequence alignments of different EMO sequences (*S. aciditrophicus*: SYN_02638; *Smithella* sp.: GX642_07725; *Syntrophomonas wolfei*: Swol_0698; *Pelotomaculum schinkii*: Psch_01459; *Syntrophorhabdus aromaticivorans*: GXY80_08670 *Geobacter metallireducens*: Gmet_2070; *Desulfosarcina ovata*: DSCOOX_18850; *M. tuberculosis*: Rv0338c; *M. smegmatis*: MSMEG_0690; *Bacillus subtilis*: BSU37180; *Haloferax volcanii*: HVO_0215; *Archaeoglobus fulgidus*: AF_0755; *Pseudomonas aeruginosa*: PA5399) were made using Clustal Omega^60^ via the EMBL-EBI framework^61^. For phylogenetic analysis of NCB domain-containing proteins, further protein sequences were included (Supplementary Tab. 2), e.g. HdrB subunits of HDRs or GlpC of anaerobic glycerol-3-phosphate dehydrogenases. A phylogenetic tree was constructed based on these Clustal Omega-aligned sequences in MEGA12^62^. The Maximum Likelihood method and a Jones-Taylor-Thornton matrix-based model^63^ were employed with a bootstrap^64^ value of 1,000. A total of 53 amino acid sequences was used, with 1,629 positions in the final dataset.

## Data availability

The protein structural data generated in this study have been deposited in the Protein Data Bank at http://www.pdb.org under the accession codes 26DL / EMD-80561 (EMO) and 26DT / EMD-80568 (EMO:ETF). The source data underlying the biochemical characterization in the main text, Fig. 2, 5, and 7, and Supplementary Fig. 1, and 6-10 are provided in the Source Data file. The mass spectrometry protein data have been deposited to the ProteomeXchange Consortium via the PRIDE^65^ partner repository with the dataset identifiers PXD068677 and PXD075953. Additional data that support the findings of this study are available from the corresponding authors upon request. Source data are provided with this paper.

## Acknowledgements

This research was supported by Deutsche Forschungsgemeinschaft (SFB 1381, project ID 403222702 to L.Z., L.A., F.D., P.H., M.B., and O.E.) and the European Research Council (Horizon Europe AdG no. 101141673 to O.E.). High-performance computing resources were available at the bwHPC cluster of the federal state of Baden-Württemberg and Deutsche Forschungsgemeinschaft (INST 35/1597-1 FUGG). The Titan Krios G4 cryo-TEM at the Cyro-EM Facility of the University of Freiburg is supported by Deutsche Forschungsgemeinschaft (project ID 506518771) and operated within the Microscopy and Image Analysis Platform (MIAP) and the Life Imaging Centre (LIC, Freiburg).

We thank Fabrice Krier for performing initial EMO-ETF interaction experiments using size-exclusion chromatography and crosslinking, Samuel Wied for preparing the plasmid encoding the two NCB domains of EMO for heterologous production in *E. coli*, and Bettina Knapp for preparing samples for LC-MS/MS and running analysis.

## Author contributions

D.K. enriched EMO and ETF from S. aciditrophicus wild-type cells, heterologously produced cyclohexanoyl-CoA DH from S. aciditrophicus in E. coli and enriched the protein, heterolo-gously produced, enriched and characterized the truncated version of EMO containing the non-cubane [4Fe:4S] clusters in E. coli, prepared and measured samples for UV/vis spectros-copy, prepared samples for EPR measurements, analysed biochemical and spectroscopic data, prepared figures, and contributed to writing of the paper. L.Z. prepared and measured samples for cryo-EM, processed data, built and refined the structural models, and contributed to writing the paper. L.A. performed experiments on protein-protein interactions between EMO and its interaction partners. L.H. measured and evaluated EPR data and contributed to writ-ing the paper. F.D. and P.H. carried out high-resolution mass spectrometric analyses, evaluat-ed data, and contributed to writing the paper. O.E. evaluated and refined data obtained dur-ing cryo-EM analysis, prepared structural models, figures, and contributed to writing the pa-per. M.B. designed the study, analysed data and contributed to writing the paper. All authors discussed the results and reviewed the manuscript.

