## Supplemental tables and figures for "Powering methanogenesis from fatty acids by a twin-heme-mediated reverse redox-loop"

**Supplementary Tables 1-2**

**Supplementary Figures 1-14**

**Supplementary Table 1 | Cryo-EM data collection, refinement, and validation statistics.**

| **Data Set** | **EMO** | **EMO:ETF** |
|---|---|---|
| PDB ID | 26DL | 26DT |
| EMBD ID | EMD-80561 | EMD-80568 |
| microscope | Titan Krios G4 | Titan Krios G4 |
| detector | Falcon 4i | Falcon 4i |
| magnification | 215,000 | 130,000 |
| voltage (kV) | 300 | 300 |
| exposure ($e^-/Å^2$) | 40 | 40 |
| defocus range (µm) | -0.6 to -1.7 | -1.0 to -2.0 |
| pixel size (Å) | 0.572 | 0.937 |
| number of movies | 10,188 | 4,885 |
| initial particles | 4,087,401 | 2,761,882 |
| imposed symmetry | $C_1$ | $C_1$ |
| final particle number | 267,857 | 306,370 |
| resolution (Å) | 1.99 | 3.0 |
| FSC threshold | 0.143 | 0.143 |
| **Refinement** | | |
| Map pixel size (Å) | 0.7627 | 0.9370 |
| Model resolution (Å) | 2.02 | 3.11 |
| FSC threshold | 0.5 | 0.5 |
| Model composition | | |
| Non-hydrogen atoms | 6,351 | 10,340 |
| Protein residues | 694 | 1,274 |
| Ligands | 20 | 21 |
| *B* factors (Å2) | | |
| Protein | 21.95 | 71.39 |
| Ligand | 41.01 | 88.70 |
| R.m.s. deviations | | |
| bond lengths (Å) | 0.004 | 0.003 |
| bond angles (°) | 0.747 | 0.649 |
| Validation | | |
| MolProbity score | 1.25 | 1.24 |
| clash score | 4.87 | 4.62 |
| poor rotamers (%) | 0.00 | 0.00 |
| Ramachandran plot | | |
| favored (%) | 98.99 | 98.58 |
| allowed (%) | 1.01 | 1.42 |
| disallowed (%) | 0.00 | 0.00 |

**Supplementary Table 2 | Proteins and genes containing NCB domains to coordinate non-cubane FeS clusters that were used for comparison and *in silico* investigations throughout this work.**

| Protein | Organism | Gene identifier | NCBI locus |
|---|---|---|---|
| **HdrB** heterodisulfide reductase subunit B | *Methanothermobacter marburgensis* | MTBMA_c04500 | CAA57038 |
| | *Methanothermobacter thermoautotrophicus* | MTH_1879 | AAB86345 |
| | *Methanothermococcus thermolithotrophicus* | A0A2D0TCB4 | WP_018154154 |
| | *Methanosarcina acetivorans* | MA_4237 | WP_011024119 |
| | *Methanospirillum hungatei* | Mhun_1837 | WP_011448821 |
| | *Desulfovibrio vulgaris* | DVU_2403 | WP_010939675 |
| | *Methanosarcina mazei* | DKM28_10735 | WP_015411490 |
| | *Archaeoglobus fulgidus* | AF_1375 | AAB89869 |
| | *Hyphomicrobium denitrificans* | Hden_0694 | WP_013214731 |
| **SdhC** succinate dehydrogenase subunit C (type E) | *Sulfolobus acidocaldarius* | Saci_0980 | WP_011277843 |
| | *Methallosphaera sedula* | Msed_0675 | WP_012020637 |
| | *Saccharolobus solfataricus* | SSO2358 | AAK42511 |
| | *Picrophilus torridus* | SAMN02745355_1129 | WP_084272933 |
| **GlcF** glycolate oxidase subunit F | *Escherichia coli* K12 | JW5486 | WP_001194661 |
| | *Shigella sonnei* | FYL34_004662 | WP_001194664 |
| | *Bacillus subtilis* | BSU28690 | WP_003229519 |
| | *Synechocystis* sp. | P73119_SYNY3 | BAA17145 |
| | *Aromatoleum aromaticum* EbN1 | ebA4492 | WP_011238356 |
| | *Pseudomonas aeruginosa* | PA5353 | WP_003114370 |
| | *Rhodobacter sphaeroides* | rsp_1018 | WP_011338662 |
| **GlpC** anaerobic glycerol-3-phosphate dehydrogenase subunit C | *Escherichia coli* K12 | JW2237 | WP_001000379 |
| | *Haemophilus influenza* | HI_0683 | NP_438843 |
| | *Shigella flexneri* | S2458 | WP_001000362 |
| | *Salmonella typhimurium* | CAC56_07820 | WP_202808681 |
| **TfrB** thiol:fumarate reductase subunit B | *Methanothermobacter marburgensis* | MTBMA_c04210 | WP_013295248 |
| | *Methanobrevibacter arboriphilus* | MBBAR_23c00060 | WP_080460913 |
| | *Methanosphaera stadtmanae* | Msp_1044 | WP_011406627 |
| | *Methanococcus maripaludis* OS7 | MMOS7_11760 | WP_119721052 |
| **HdrD** heterodisulfide reductase subunit D | *Methanosarcina barkeri* | Mbar_A1599 | CAA70997 |
| | *Methanosarcina thermophila* | MSTHT_2244 | WP_048167969 |
| | *Methanosarcina mazei* | MM_1844 | WP_048045886 |
| | *Methanosarcina acetivorans* | MA_0688 | WP_011020733 |
| **BamD** class II benzoyl-CoA reductase subunit D | *Geobacter metallireducens* | Gmet_2085 | ABB32314 |
| | *Syntrophus aciditrophicus* | SYN_00259 | WP_011416388 |
| | *Desulfosarcina cetonica* | - | WP_213182121 |
| | *Desulfococcus multivorans* | dsmv_2374 | WP_020876805 |
| **EMO(-like)** ETF:(M)MK oxidoreductase | *Syntrophus aciditrophicus* | SYN_02638 | WP_011418540 |
| | *Smithella* sp. | GX642_07725 | NLD81036.1 |
| | *Syntrophomonas wolfei* | Swol_0698 | WP_011640127 |
| | *Geobacter metallireducens* | Gmet_2070 | WP_004514033 |
| | *Desulfosarcina ovata* | DSCOOX_18850 | WP_155309984 |
| | *Mycobacterium tuberculosis* | Rv0338c | WP_003916653 |
| | *Mycobacterium smegmatis* | MSMEG_0690 | WP_011727122 |
| | *Bacillus subtilis* | BSU37180 | WP_003244084 |
| | *Haloferax volcanii* | FQA18_14375 | WP_004065535 |
| | *Archaeoglobus fulgidus* | XD40_1303 | KUJ93526 |
| | *Pseudomonas aeruginosa* | PA5399 | NP_254086 |
| | *Pelotomaculum schinkii* | Psch_01459 | WP_190239682 |
| | *Syntrophorhabdus aromaticivorans* | GXY80_08670 | NLW35536 |
| **D-iLDH** NADH-independent D-lactate dehydrogenase | *Pseudomonas putida* | PP_4737 | WP_010955373 |
| | *Pseudomonas aeruginosa* | PA4772 | WP_003123510 |
| | *Shewanella oneidensis* | SO_1521 | WP_011071700 |
| | *Vibrio cholerae* | VC0395_0253 | WP_000188686 |

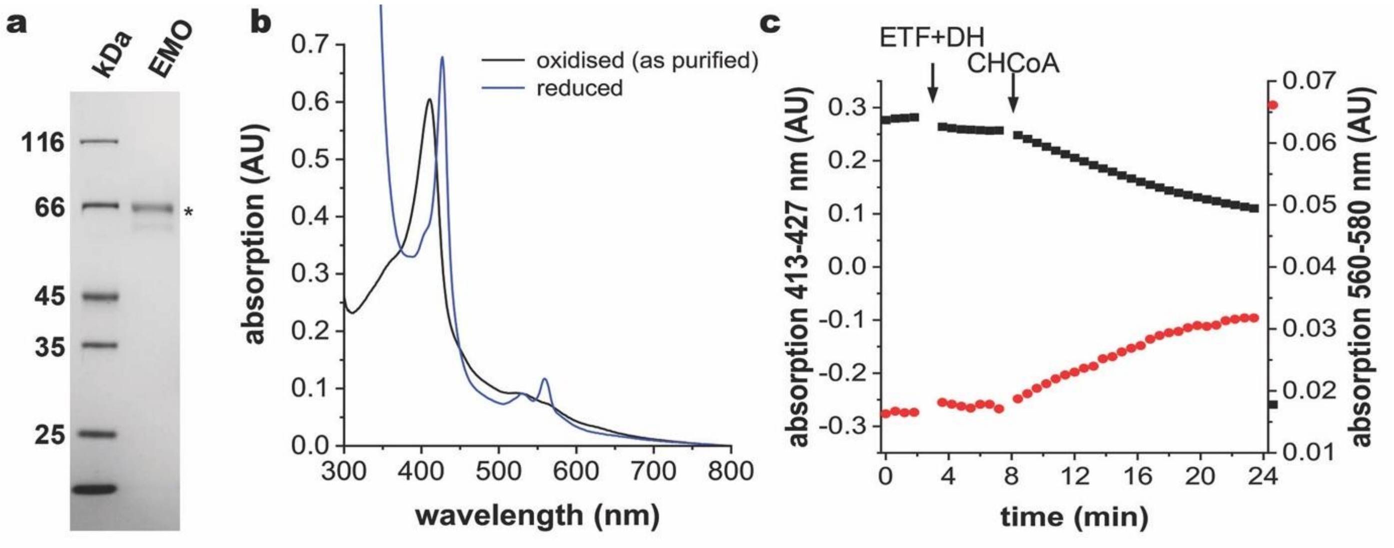


**Supplementary Figure 1 | Enrichment of EMO from *S. aciditrophicus* wild-type cells and electron transfer activity of EMO.** **a**, SDS-PAGE analysis of enriched EMO (3 µg). The protein band marked with an asterisk was excised and analysed by UPLC-MS, resulting in a 30.6 % sequence coverage for EMO (SYN_02638) as the single hit. **b**, UV/vis absorption spectra of oxidised (as purified) EMO (3 µM) and fully dithionite-reduced EMO. **c**, Time-dependent heme *b* cofactor reduction of EMO (3 µM) by cyclohexanoyl-CoA (CHCoA, 100 µM) in the presence of CHCoA DH (2 µM) and ETF (6 µM). The differences in absorption at 413 and 427 nm (Soret band, black squares) and between 560 and 580 nm (red circles) are shown.

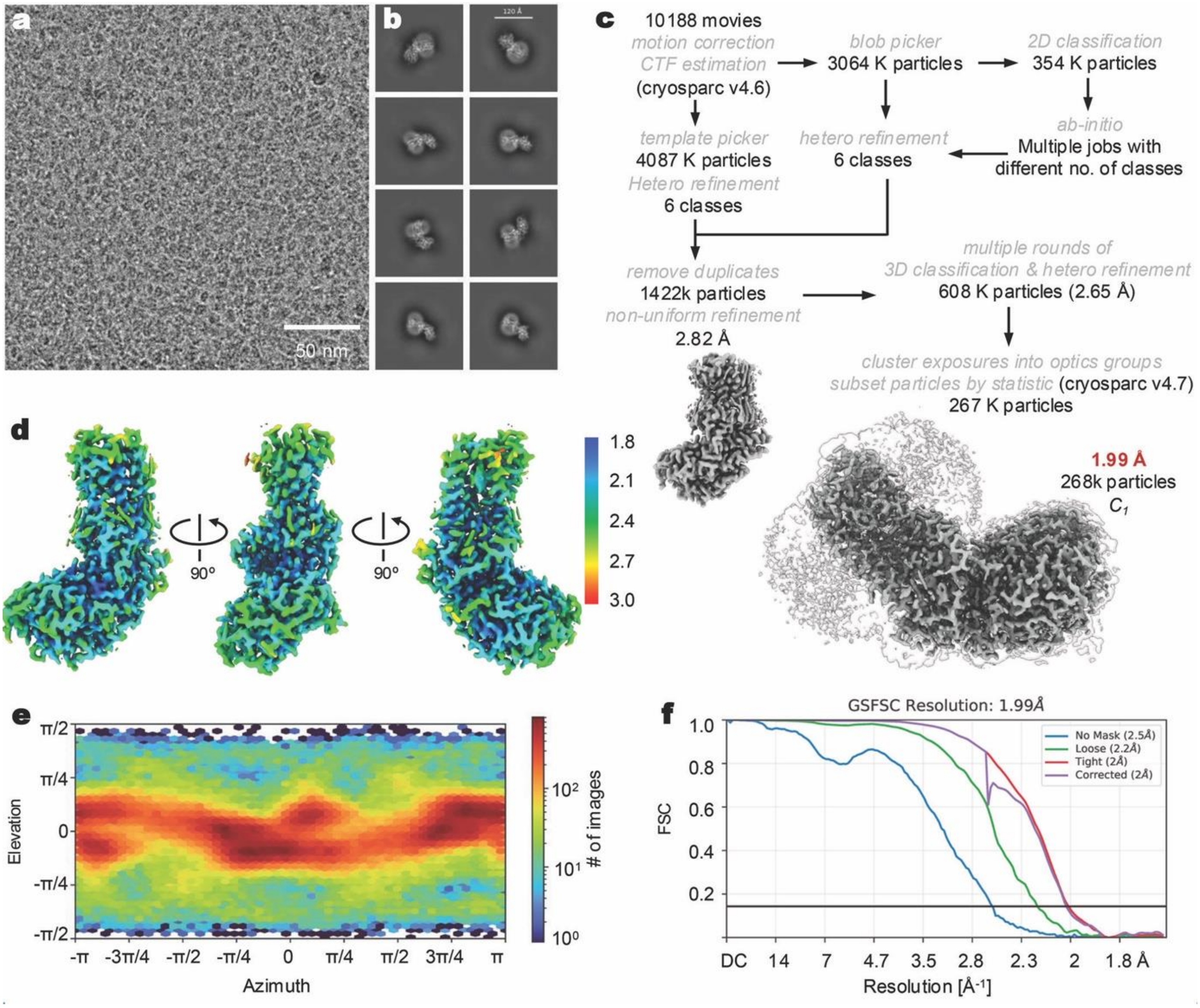


**Supplementary Figure 2 | Data statistics and workflow for the cryo-EM data processing of EMO.** **a**, Representative micrograph from 10,188 movies. **b**, 2D class averages with different views. **c**, Workflow for the data processing, leading to a refined map at 2.0 Å resolution using 268 K particles. **d**, Local resolution maps for different orientations of EMO. **e**, Direction distribution of particles used for 3D reconstruction. **f**, Fourier shell correlation (FSC) curves with a gold standard cutoff of 0.143.

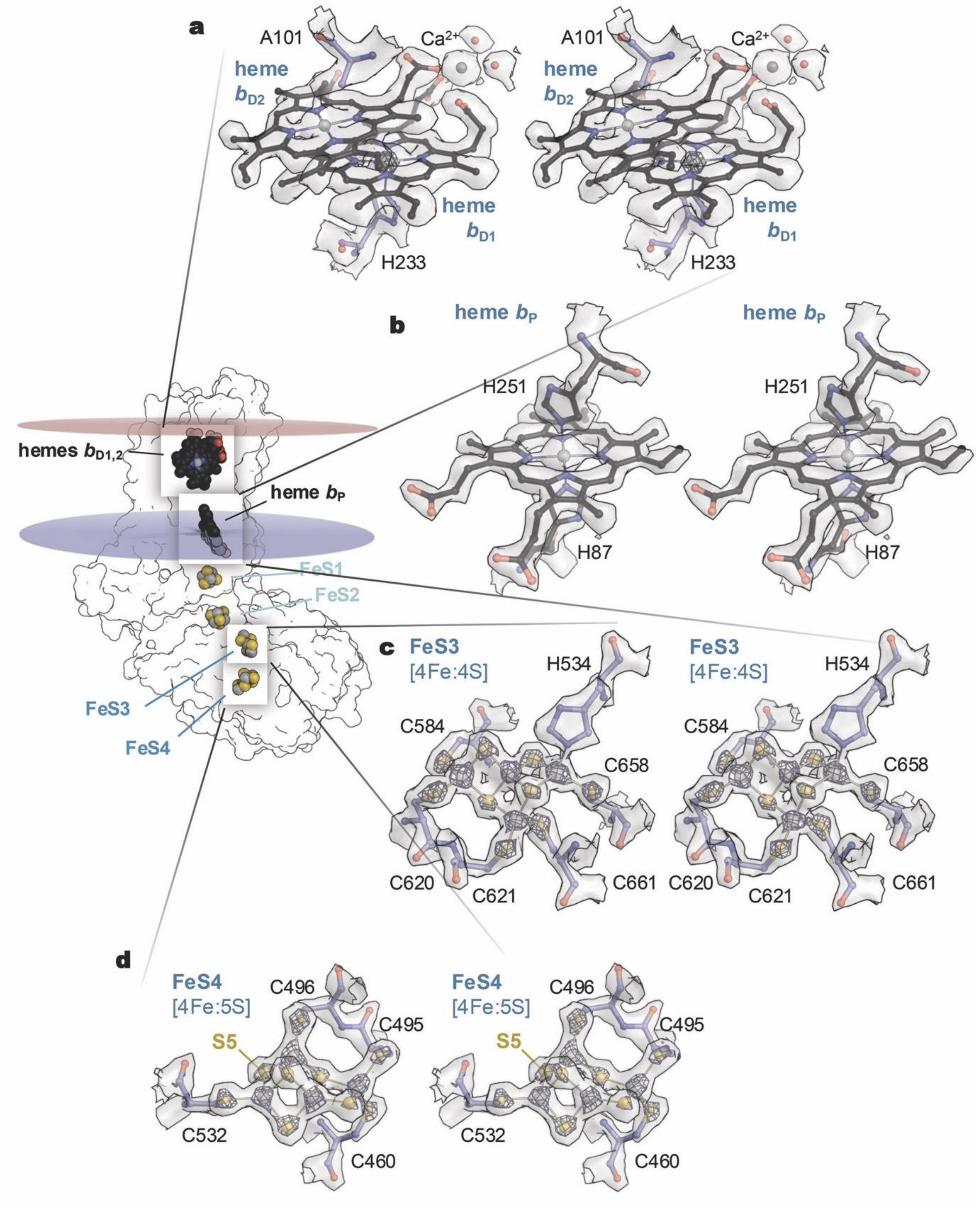


**Supplementary Figure 3 | Representative electron density maps showing selected cofactors of EMO.** **a**, The two stacked distal heme *b* groups, $b_{D1}$ and $b_{D2}$. Note the additional density feature at the axial position of heme $b_{D2}$, near residue A101. **b**, The proximal, *bis*-histidine-coordinated heme $b_P$. **c**, The iron-sulfure cluster FeS3, a typical non-cubane [4Fe:4S] cluster. **d**, FeS4, the last cluster in the chain of cofactors, with an additional sulfide, S5, making it a novel [4Fe:5S] moiety. All sites are shown as wall-eyed stereo renderings.

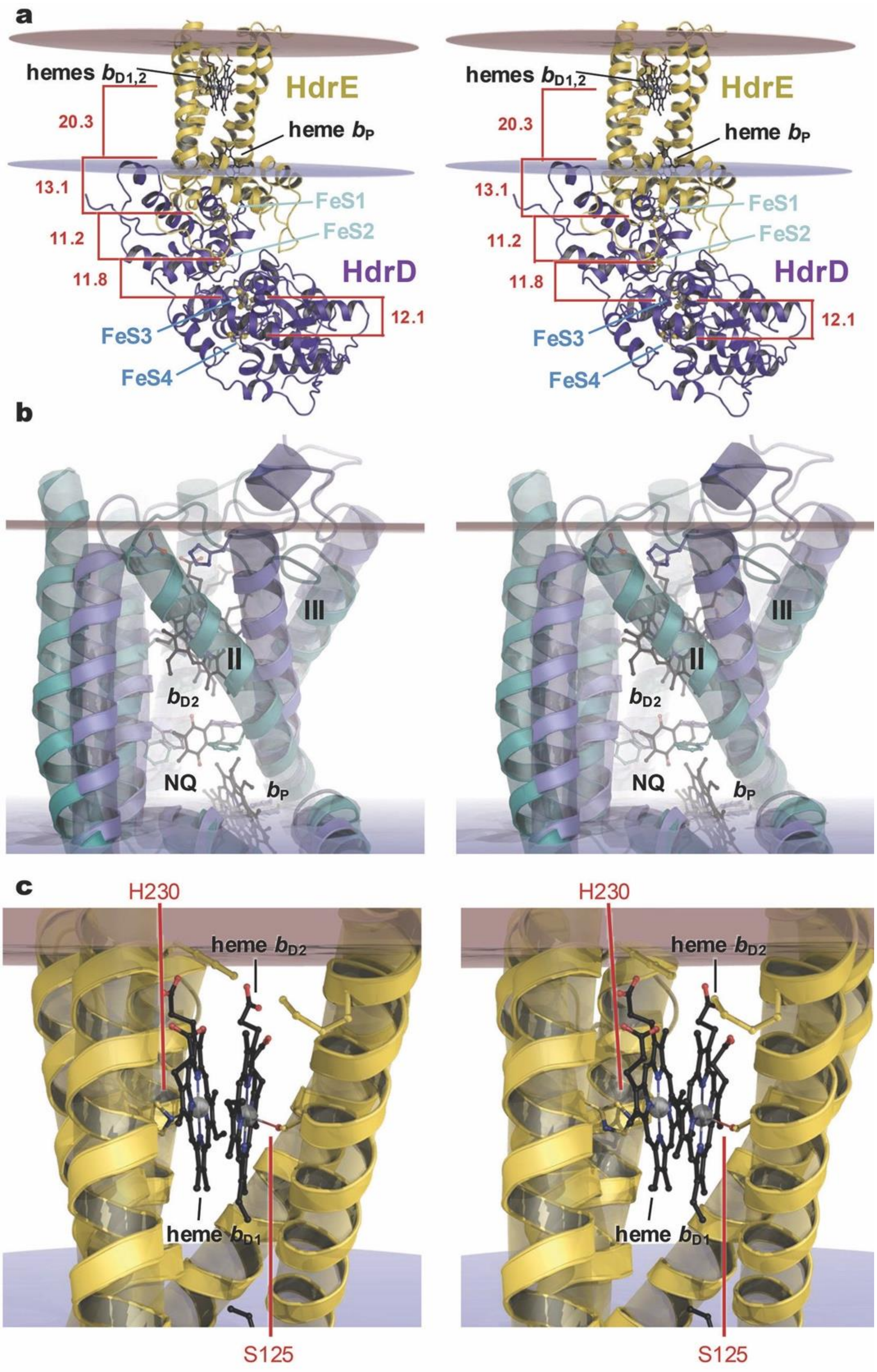


**Supplementary Figure 4 | Structural comparison of membrane-bound heterodisulfide reductase HdrDE, *Ms*EMO and *Sa*EMO.** **a**, AlphaFold model of MaHdrDE placed in a membrane. Numbers denote inter-cofactor distances. The prediction included three heme *b* groups that were placed precisely as seen in *Sa*EMO, with a stacked pair $b_{D1,2}$. **b**, Structural differences between the TM domains of *Sa*EMO (blue) and *Ms*EMO (cyan, PDB 9OJN). In the latter, TM helix II is substantially tilted, blocking the binding site for the two distal heme groups. The bound naphtoquinone (NQ) modelled in *Ms*EMO could not be accommodated in *Sa*EMO. **c**, Detail of the distal heme stack of the *Ma*HdrDE model (a). Heme $b_{D1}$ is coordinated by H230, while heme $b_{D2}$ has S125 as an axial ligand.

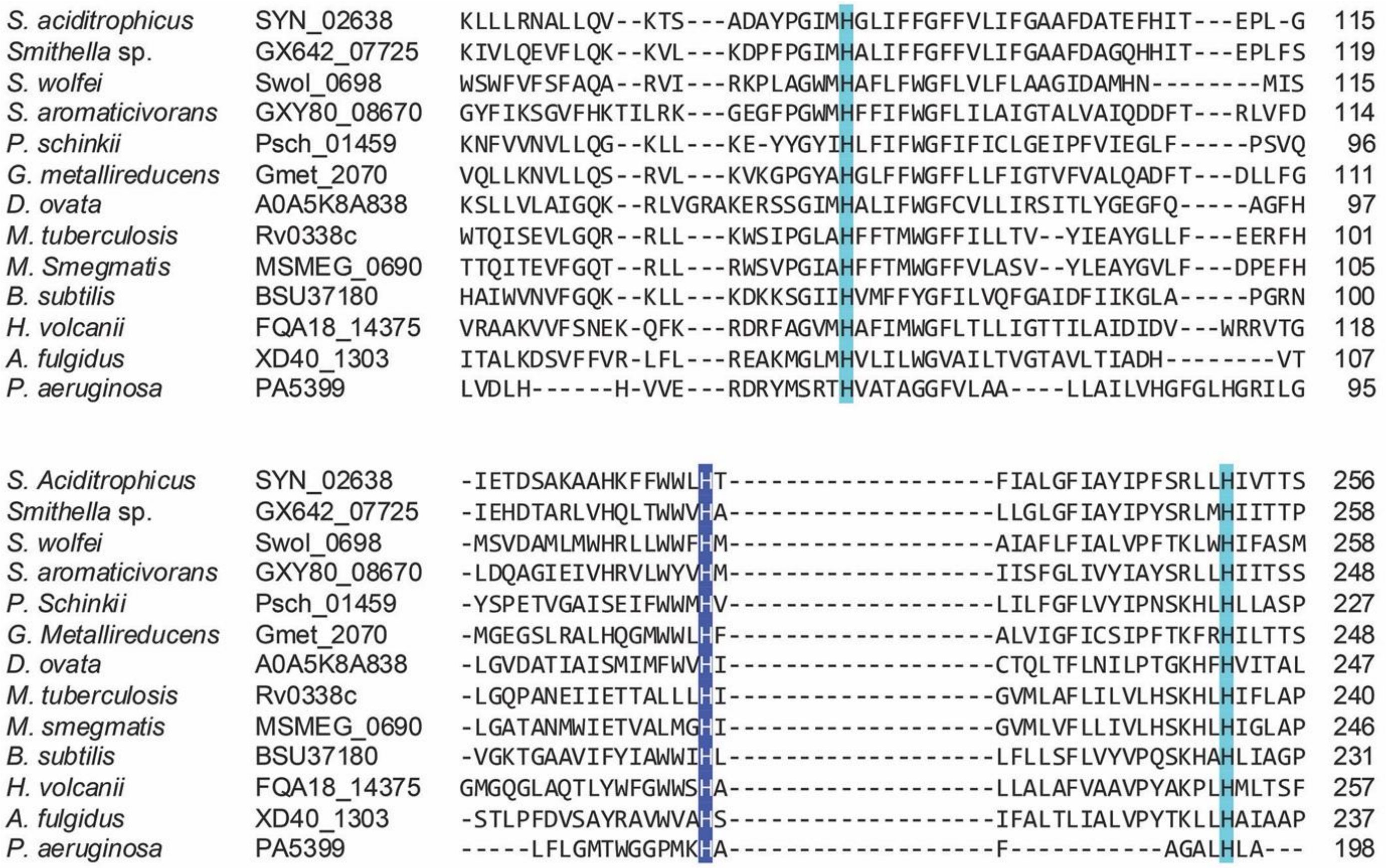


**Supplementary Figure 5 | Multiple sequence alignment of conserved histidine residues.** These residues potentially serve as axial heme *b* ligands in different bacterial and archaeal *emo* genes (dark blue: distal heme $b_D$; light blue: proximal heme $b_P$). Note that the gene products of *M. tuberculosis* (Rv0338c) and *M. smegmatis* (MSMEG_0690) are also referred to as EtfD, of *B. subtilis* (BSU37180) as FadF and of *P. aeruginosa* as dimethylglycine demethylation protein DgcB.

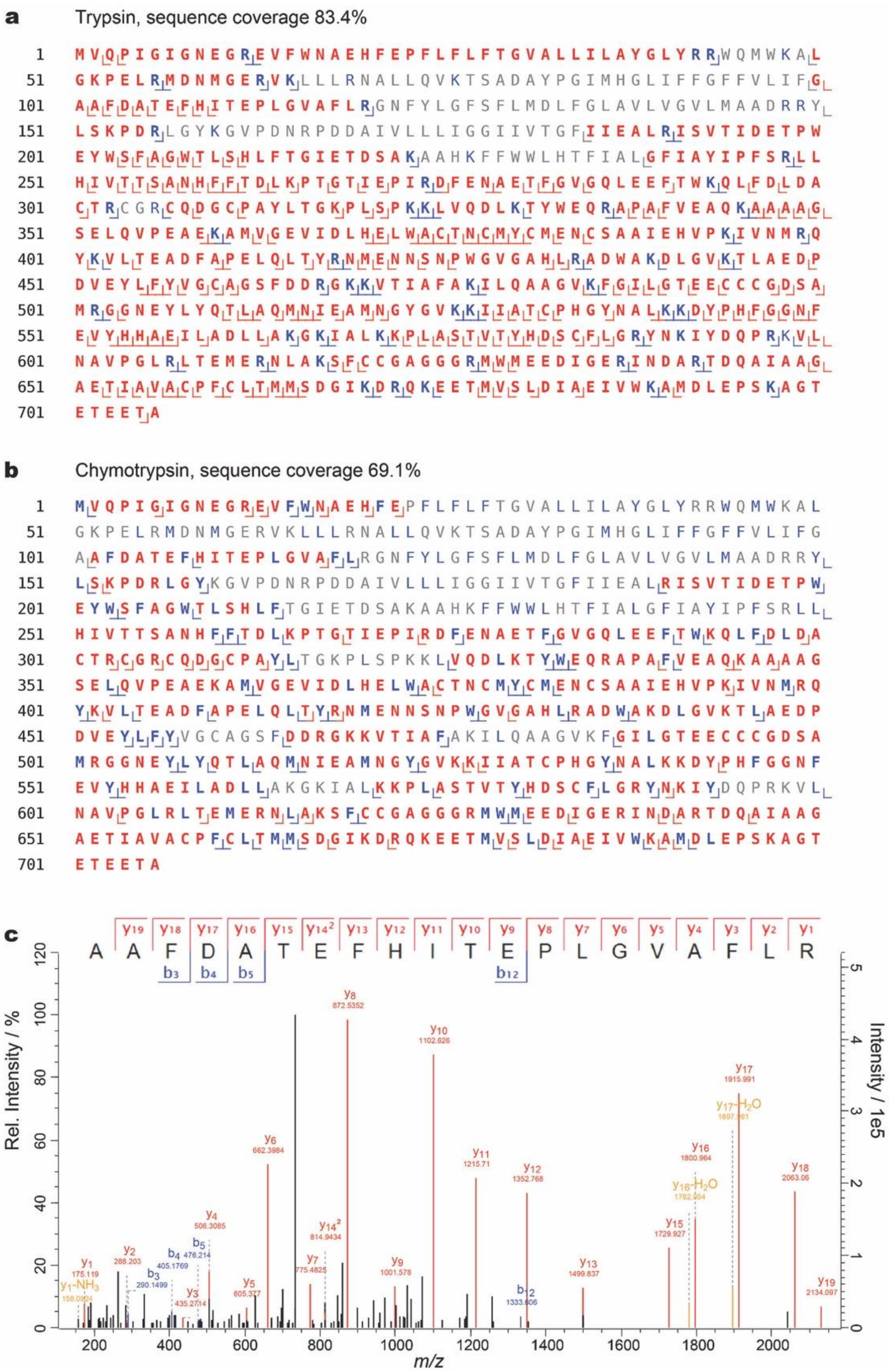


**Supplementary Figure 6 | LC MS/MS analysis. The data shows no evidence of post-translational modification of A101 to explain the observed density maximum near the stacked hemes $b_{D1,2}$ (Supplementary Fig. 3a).** **a**, Sequence coverage for EMO after tryptic digest, and **b**, digestion with chymotrypsin. Amino acids covered by identified peptides are shown in bold red and bold blue, uncovered regions in grey. Blue denotes amino acids preceding regular proteolytic cleavage sites. Positions of observed cleavages are additionally indicated below the sequence. **c**, Representative MS/MS spectrum identifying semi-tryptic peptide A101-R120, with all matching *b* (blue) and *y* ions (red) indicated above the corresponding peaks and in the inset scheme. Results were obtained by data-dependent acquisition and confirmed using data-independent acquisition mode.

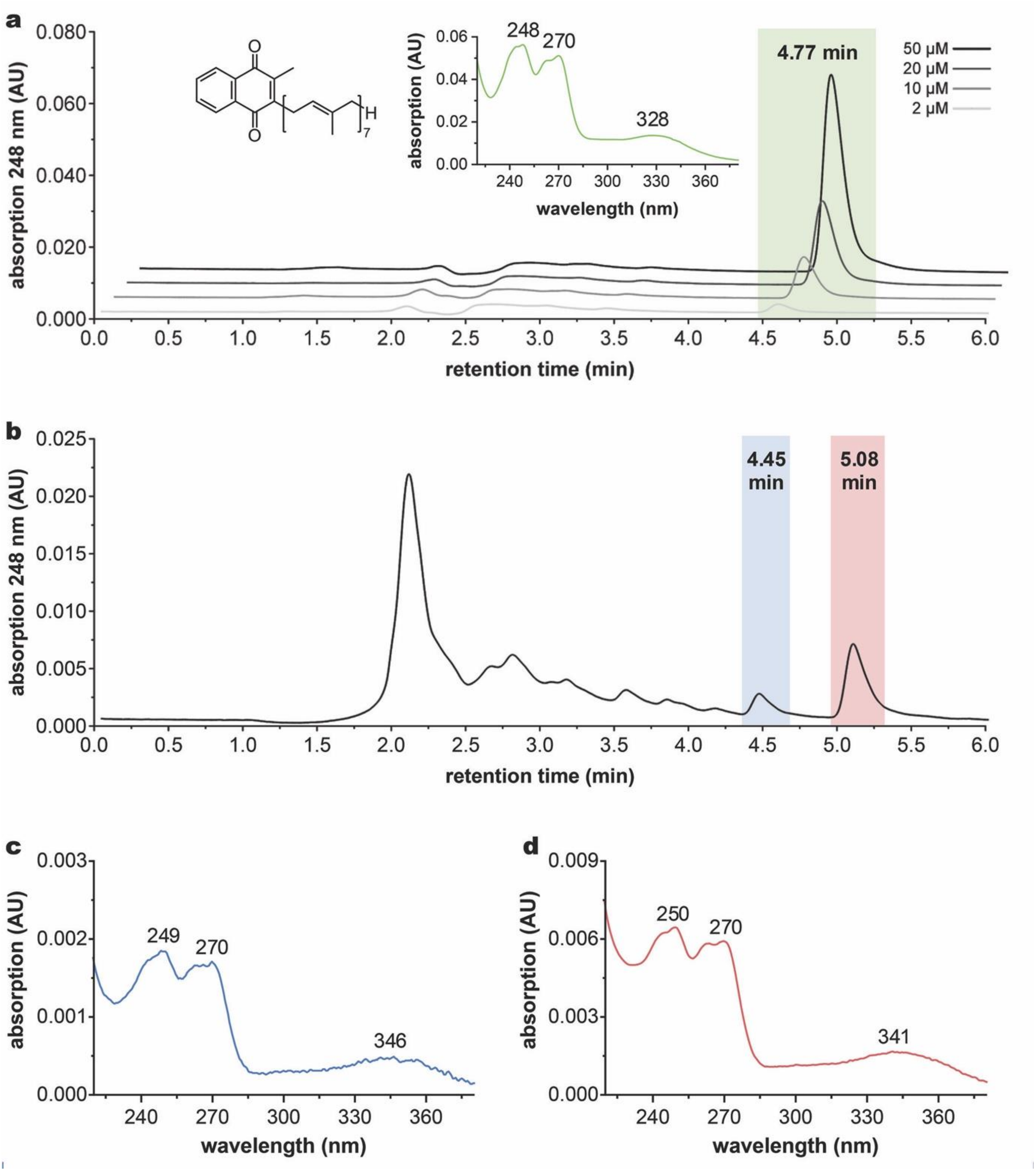


**Supplementary Figure 7 | UPLC analysis of MMK species extracted from enriched *Sa*EMO. a**, UPLC chromatogram of authentic MK-7 standards at 248 nm, eluting around 4.8 min with the molecule structure and UV spectrum shown as insets. **b**, UPLC chromatogram at 248 nm of an ethanol treated EMO sample. Only the two highlighted peaks exhibited typical spectra of MMK species (**c**, 4.45 min peak and **d**, 5.08 min peak).

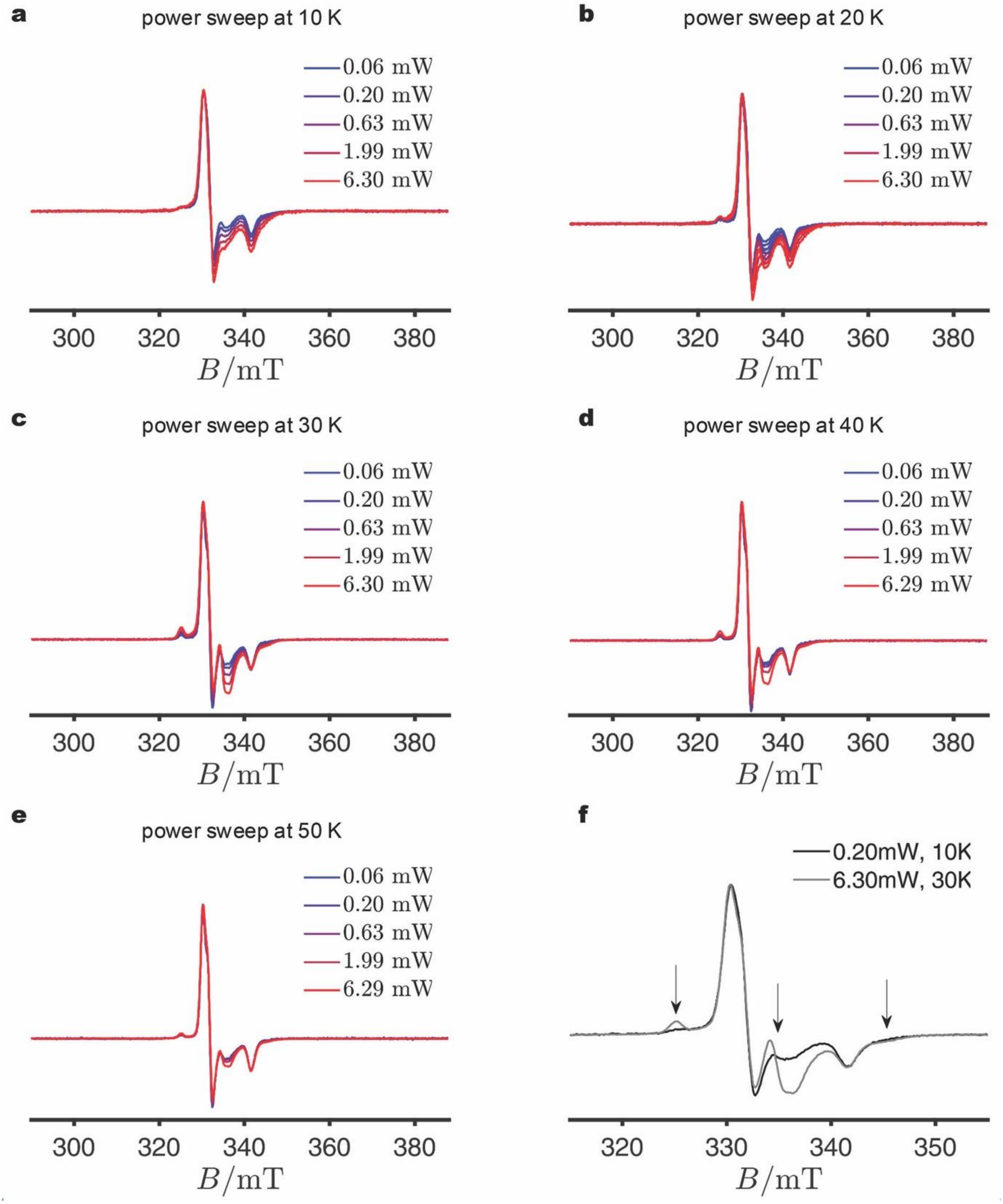


**Supplementary Figure 8 | Power and temperature dependence of the *Sa*EMO NCB domain during EPR spectroscopy. a-e**, Power sweeps recorded at 10, 20, 30, 40, and 50 K. **f**, Comparison of EPR spectra under the indicated conditions, highlighting the distinct power- and temperature-dependent behaviours of the two non-cubane clusters. Individual spectral features cannot yet be assigned to specific clusters.

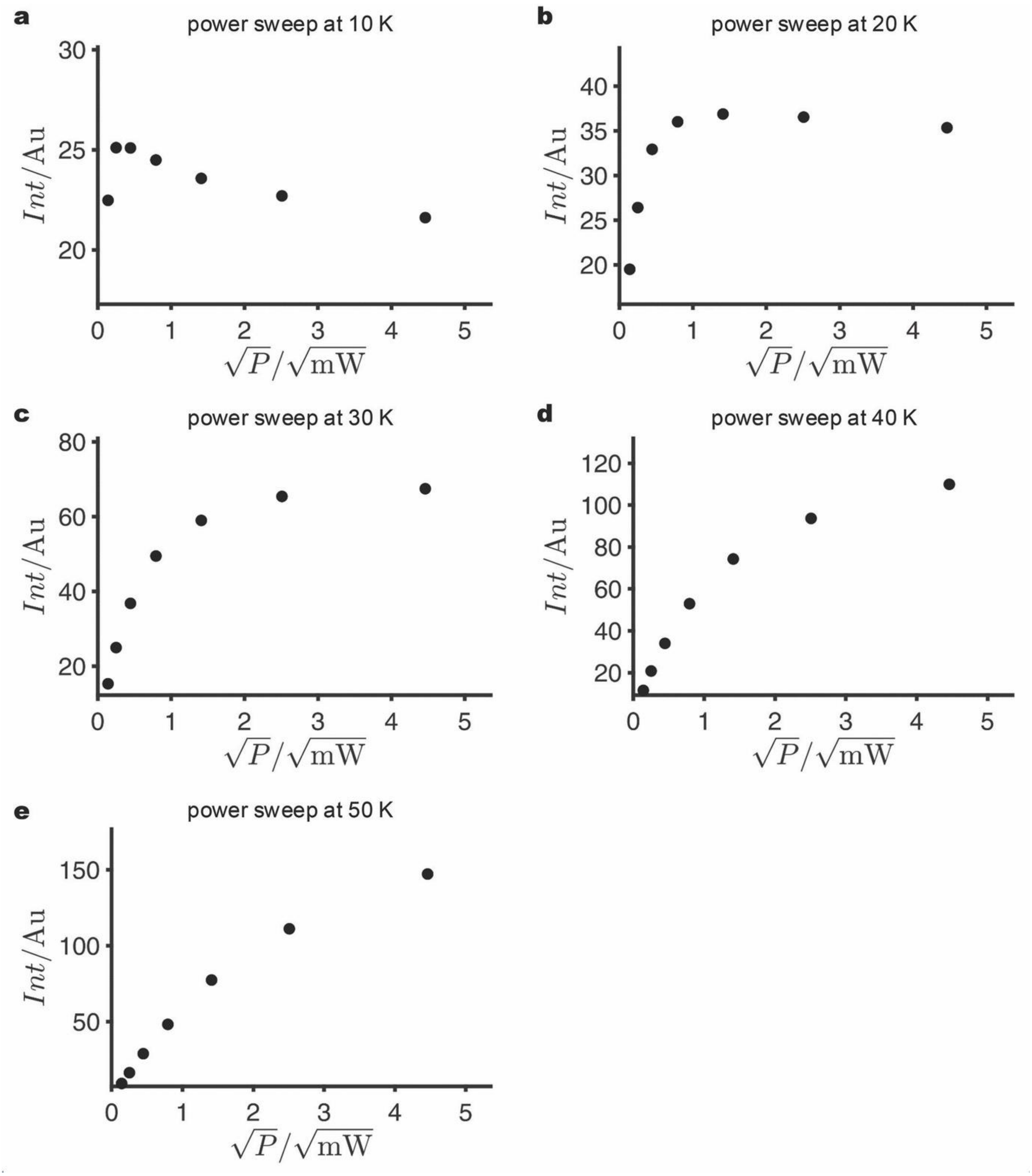


**Supplementary Figure 9 | Power saturation behaviour of the *Sa*EMO NCB domain during EPR spectroscopy. a-e**, Power sweeps recorded at 10, 20, 30, 40, and 50 K.

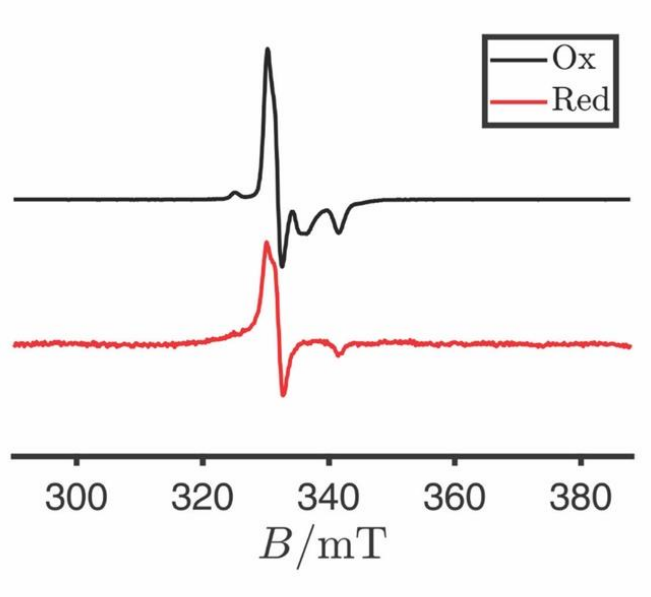


**Supplementary Figure 10 | EPR spectra of the *Sa*EMO NCB domains.** Expanded view of the reduced spectrum shown in Fig. 5e, highlighting complete signal bleaching of FeS cluster species 2 in this sample.

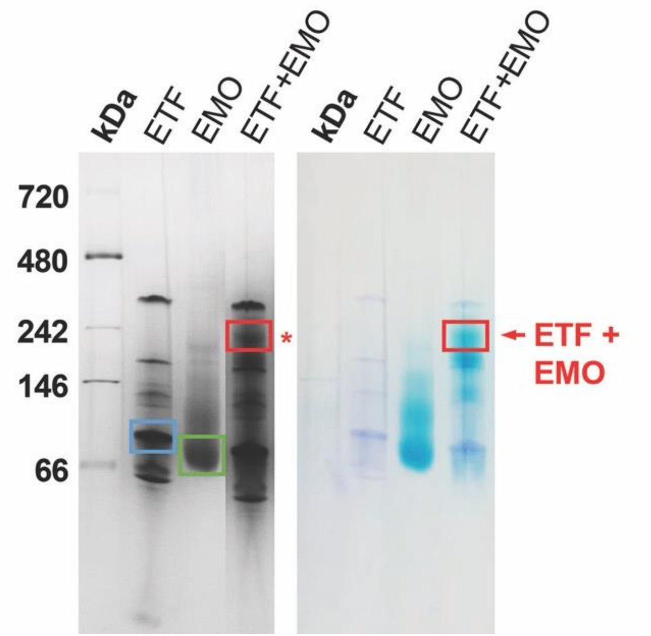


**Supplementary Figure 11 | Blue Native-PAGE analysis showing complex formation of EMO and ETF from *S. aciditrophicus*.** Left, Coomassie-stained 4–16% native gel; right, in-gel heme staining of the same gel. EMO and ETF were loaded at 10 µg each. For the EMO–ETF mixture, EMO was 10 µg and ETF adjusted to a 1:1 molar ratio, accounting for enrichment impurities. The band marked with a red star was excised and analyzed by UPLC–MS, yielding sequence coverages of EtfA 57.9%, EtfB 54.3%, and EMO 24.7%.

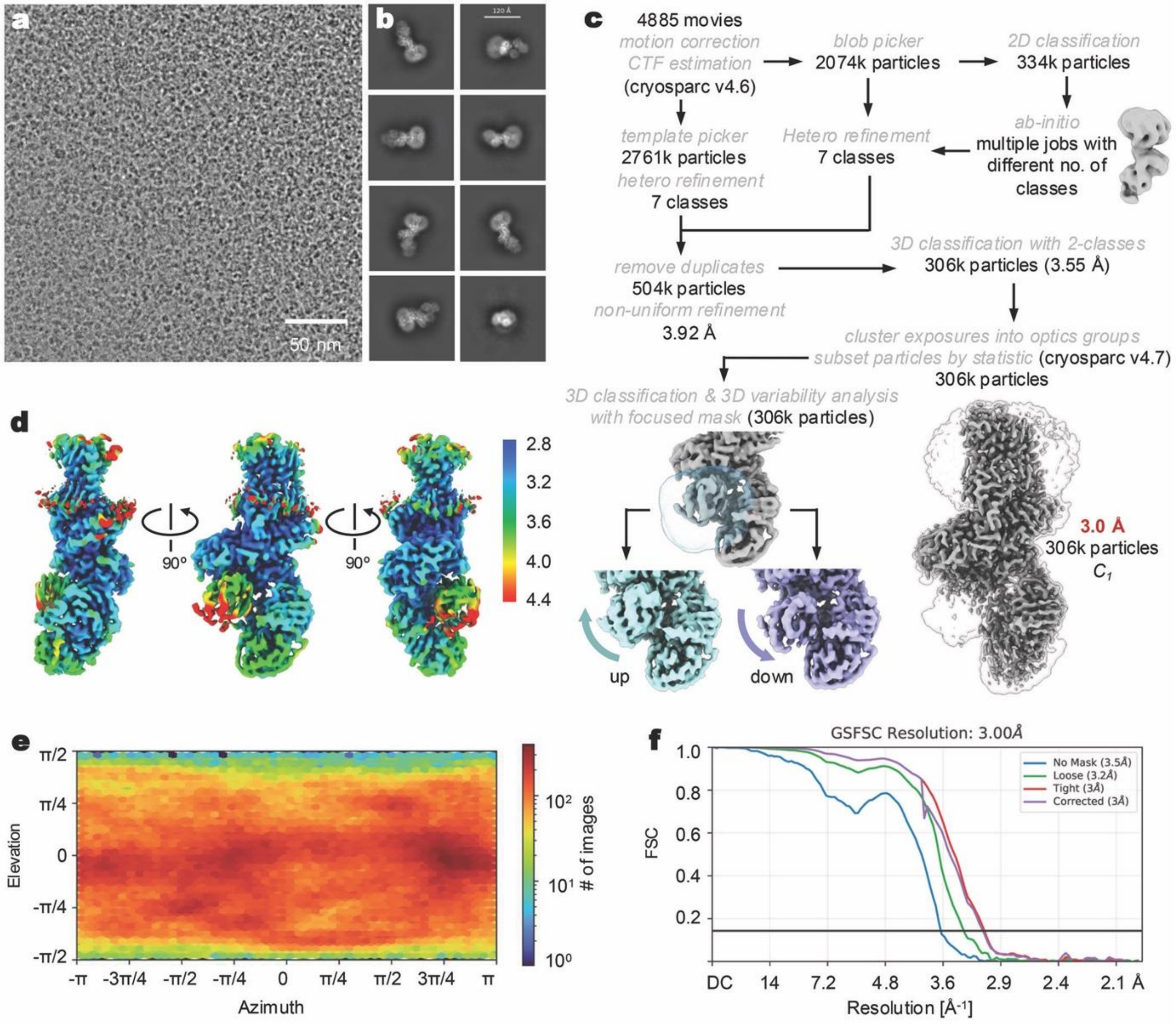


**Supplementary Figure 12 | Data statistic and workflow for the cryo-EM data processing of the EMO-ETF complex. a**, Representative micrograph of 4,885 movies. **b**, 2D class averages with different views showed the binding of ETF to EMO. **c**, Workflow for the data processing, and the final resolution was at 3.0 Å using 306 K particles. 3D classification and 3D variability analysis showed the conformational changes of the FAD-binding domain of ETF. **d**, Local resolution maps for different orientations of the EMO-ETF complex. **e**, Direction distribution of particles used for 3D reconstruction. **f**, Fourier shell correlation (FSC) curves.

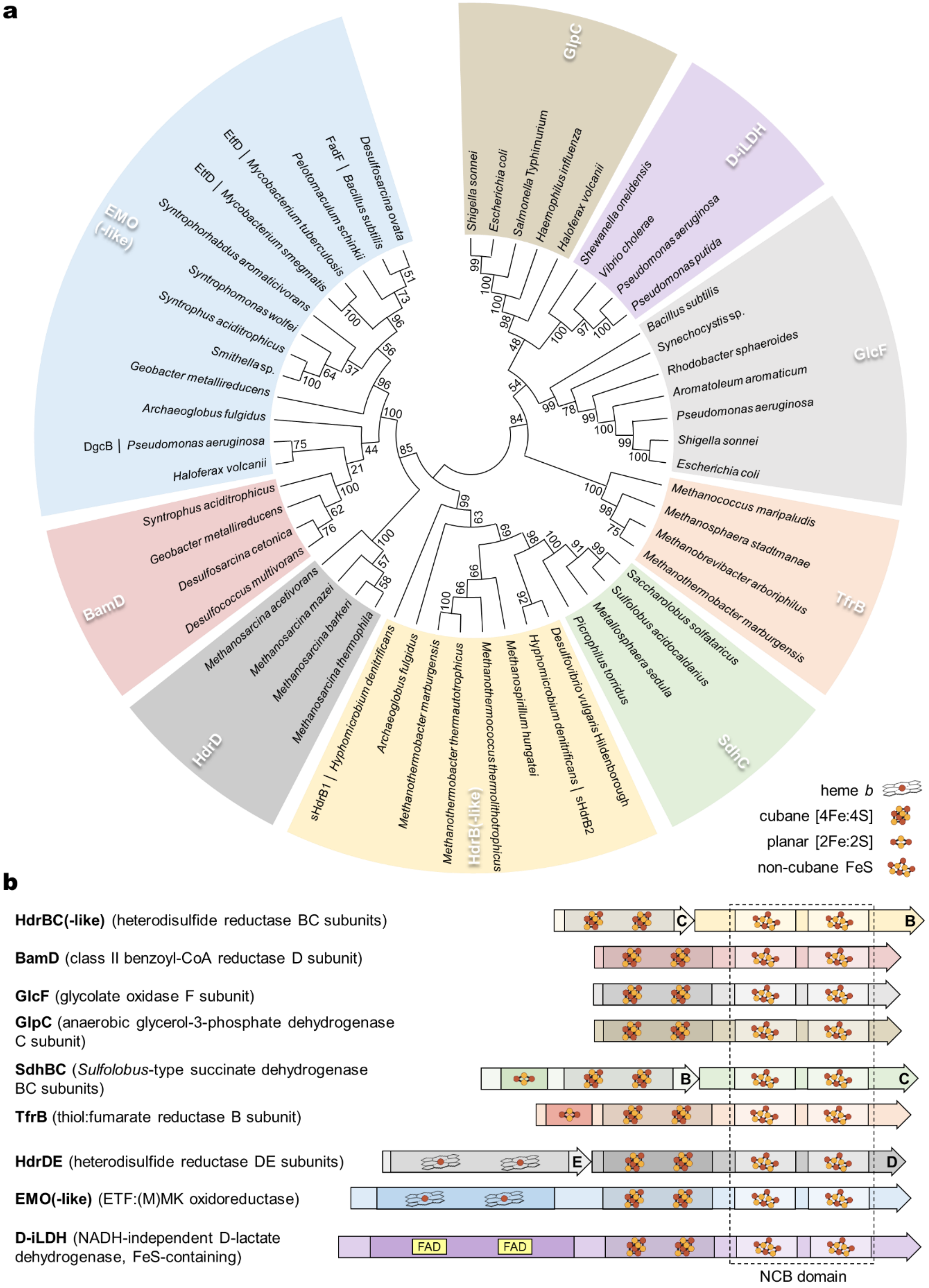


**Supplementary Figure 13 | Phylogenetic analysis of proteins harbouring NCB domains (formerly cysteine-rich CCG domains). a**, Unrooted phylogenetic tree of NCB domains, highlighting distinct clades. The tree was inferred using the Maximum Likelihood method with the Jones-Taylor-Thornton model and 1,000 bootstrap values. **b**, Schematic domain architectures of representative NCB-containing enzymes. Corresponding gene accession numbers are provided in Supplementary Table 2.

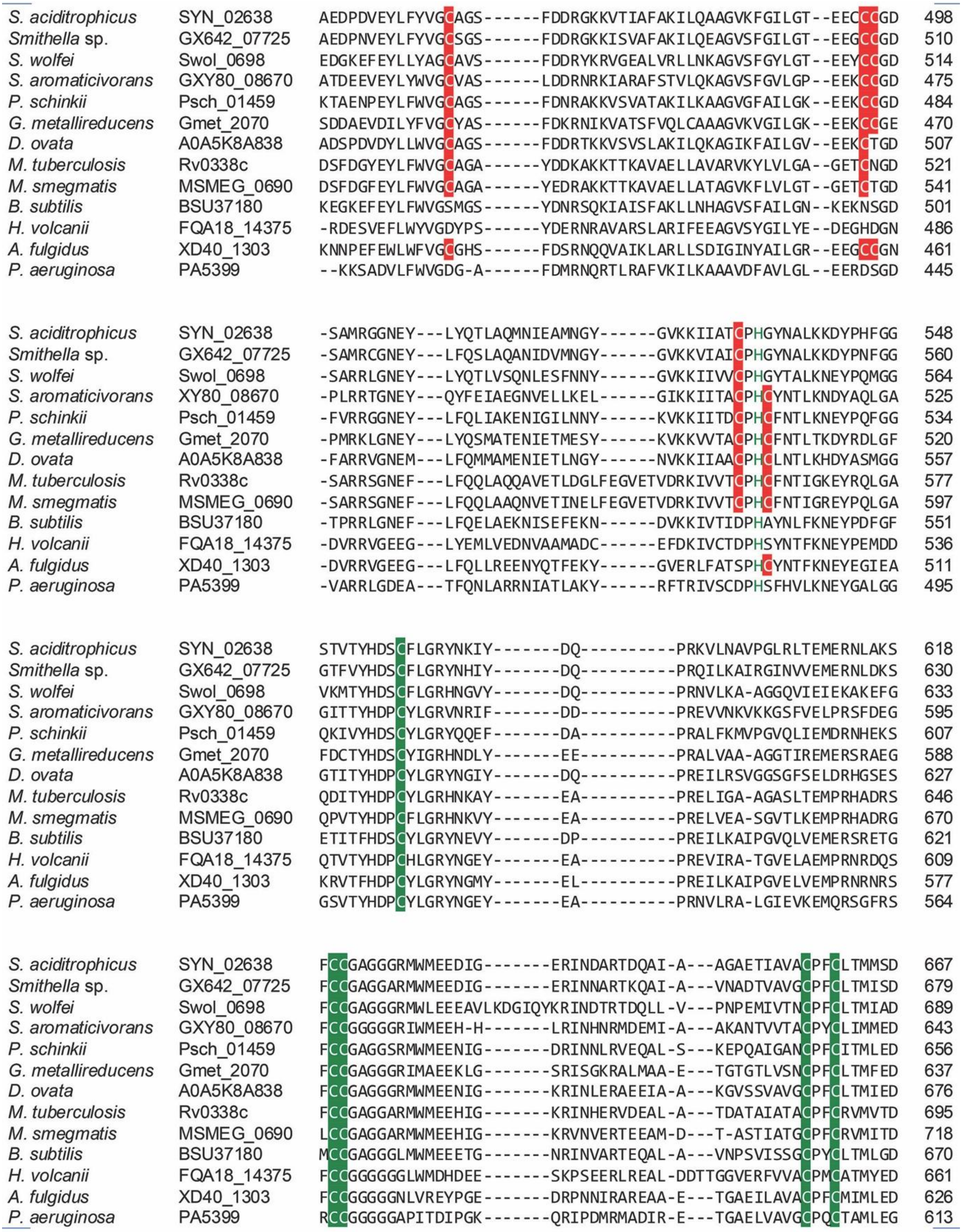


**Supplementary Figure 14 | Sequence alignment of NCB domains highlighting non-cubane cluster-binding residues.** Residues coordinating the distal FeS4 cluster are highlighted in red, those coordinating the distal FeS3 cluster in green.
